# Phenotypic inference from sparse tumor genomes informs an explainable deep-learning model for cancer prognosis

**DOI:** 10.64898/2026.06.26.734894

**Authors:** Sydney Grant, Aritro Nath

**Affiliations:** Department of Medical Oncology & Therapeutics Research, City of Hope

## Abstract

Somatic genomic alterations are widely profiled in cancer and remain the primary source for personalized therapy, yet their clinical utility is limited to few actionable targets. AI/ML models offer opportunities to capture genome-wide complexities, but clinical translation is hindered by poor interpretability, often limited to single-gene effects, and overlooks higher-order phenotypic interactions. To address this, we developed PhenoMap, a machine-learning framework that infers tumor phenotypic states from somatic variants. Trained on 9,000 pan-cancer genomes and transcriptomes, PhenoMap accurately reconstructs expression-based pathway enrichment scores and consolidated hallmark cancer phenotypes, enabling multilevel interpretation at phenotype, pathway, and gene scales. PhenoMap captured molecular subtypes and key resistance pathways across breast, lung, and brain cancers. We leveraged these features in PhenoSurv, a deep survival model integrating phenotypic reconstruction loss, Kullback–Leibler divergence, and survival loss to learn biologically-grounded predictors. PhenoSurv outperformed state-of-the-art survival models while providing robust mechanistic explanations. *NOTCH1* signaling and *SMARCA4* mutations emerged as a major prognostic factor in hormone receptor–positive breast cancer. TGF-β signaling and inflammasomes, potentially modulated by *FAT1*, predicted lung adenocarcinoma outcomes, while inositol metabolism and PI3K signaling were key drivers in brain cancer. Together, PhenoMap and PhenoSurv provide accurate, interpretable, and clinically actionable models for precision oncology.

**Graphical Abstract:** 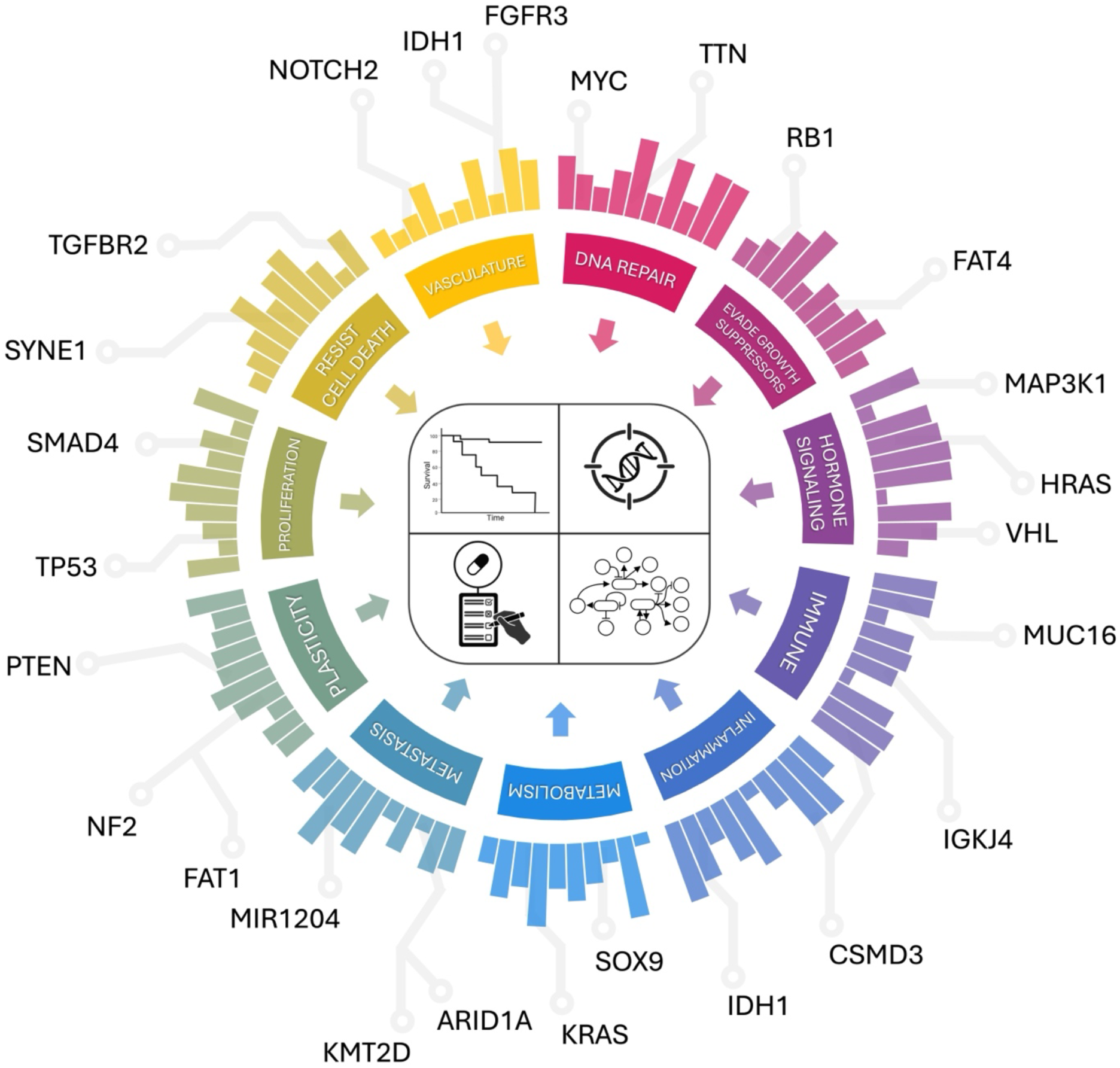

PhenoMap framework leverages genomic data and explainable deep learning to identify phenotype, pathway, and gene-level prognostic markers for precision oncology.

## INTRODUCTION

Precision oncology aims to tailor treatment strategies based on the molecular characteristics of individual tumors to improve patient outcomes^1^. The widespread adoption of next-generation sequencing (NGS) has enabled clinicians to identify somatic mutations^2,3^ and guide treatment decisions^4^, making genomic testing a cornerstone of cancer precision medicine^5–7^. Despite its routine use, the clinical utility of NGS testing is limited and significantly benefits only a subset of patients^8^, as only a small subset of genes, like as *BCR-ABL, EGFR, KRAS, BRAF, PIK3CA, IDH1/2,* are well-characterized for therapeutic relevance^9,10^. Most alterations detected by NGS tests are either passenger mutations or variants of unknown significance^11^, offering limited actionable insights for most patients.

To overcome these limitations, artificial intelligence and machine learning (AI/ML) models have been developed to extract predictive value from comprehensive representations of tumor genomic data, including variants in the protein-coding and non-coding regions^12^. While these AI/ML methods capture complex mutation patterns across the somatic mutation landscape, their lack of interpretability beyond individual-gene effects and lack of phenotypic-level biological relevance limit their clinical utility. Existing methods for interpreting the contribution of somatic mutations in ML models are predominantly limited to single-gene effects^13,14^. Moreover, models based on gene expression data consistently outperform those relying on somatic mutations^15^ and offer greater interpretability through pathway-level features such as those derived from gene set enrichment analysis^16,17^. In contrast, methods based on genomic alterations are less common and often lack granularity or scalability^18^. Somatic genomic alterations are a key component of clinical diagnostics and are increasingly leveraged for diagnosis, prognosis, and treatment selection. However, interpretation and prevalence of variants of unknown significance remain challenging and have led to a lack of uniformity in clinical decision making and underutilization of genomic test results^4,19^. This has highlighted the need for integrative models that can transform high-dimensional genomic data into biologically interpretable and clinically actionable insights.

In this work, we introduce a two-stage framework designed to bridge the gap between the abundance of clinical NGS data and the biological interpretability needed for precision oncology. The first stage, PhenoMap, learns an interpretable representation of somatic mutations by predicting gene expression–derived pathway activity scores from paired genomic and transcriptomic data. These pathway-level features not only capture known cancer subtypes but also reveal mechanisms such as PI3K/AKT signaling associated with therapeutic resistance.

In the second stage, these interpretable features serve as input to PhenoSurv, a deep learning model for survival prediction, enabling predictive modeling grounded in biologically meaningful phenotypes. PhenoSurv utilizes variational autoencoders (VAEs) to model survival outcomes from high-dimensional genomic features. VAEs effectively reduce data complexity by encoding somatic mutation-derived pathway scores into a latent space, enabling reconstruction with high fidelity and preserving biological relevance. This approach addresses the limitations of traditional models in handling large-scale NGS data, while enhancing predictive accuracy and interpretability in survival analysis. Prior studies, primarily using gene expression or methylation data, have shown that VAE-derived latent representations can support downstream tasks such as cancer subtyping and survival prediction^20–24^. However, standard VAEs are typically trained to minimize reconstruction loss and Kullback-Leibler divergence, which may not yield latent spaces that are biologically informative or aligned with specific clinical tasks, such as the prediction of survival outcomes. The PhenoSurv model incorporates supervision from the downstream objective of predicting disease-free survival during training to improve the relevance and interpretability of these learned representations^21^.

Together, these developments position the PhenoMap–PhenoSurv framework as a substantial advance in interpretable cancer genomics. Computationally, PhenoSurv outperformed existing deep learning survival models while preserving full interpretability, enabling survival-associated features to be traced directly to phenotype-, pathway-, and gene-level representations. This integration of biologically grounded latent features with survival modeling provides both improved predictive performance and mechanistic transparency. Biologically, downstream interpretation of PhenoSurv revealed multiple understudied pathways with strong prognostic significance across cancers, including *NOTCH1* signaling as a top predictor of survival in HR+/HER2– breast cancer, the *CLEC7A*-linked inflammasome pathway in lung cancer, and several metabolic pathways associated with outcomes in brain cancers. These findings demonstrate that our framework not only enhances the analytical power of somatic genomics but also uncovers meaningful biology that is obscured by single-gene approaches. Overall, PhenoMap and PhenoSurv together support more biologically informed and clinically actionable precision oncology strategies through interpretable deep learning.

## RESULTS

### PhenoMap overview: mapping tumor genomics to pathway activities and cancer hallmarks

Deep learning models trained on tumor genomic information often lack interpretability beyond single-gene effects, limiting their ability to provide biologically meaningful insights. In contrast, models leveraging gene expression data offer richer interpretability, such as sample-level pathway activity, but gene expression profiles are rarely collected in clinical settings. To address this gap and construct a biologically informative portrait of each tumor, we derived sample-specific pathway activity scores from somatic single-nucleotide variants (SNVs) and copy number alterations (CNAs) across 9,276 tumors spanning 33 cancer types. This transformation reflects the functional impact of genomic alterations at the pathway level and can be further condensed into hallmark cancer phenotypes and gene-level contributors, providing a compact, multi-layered, and biologically interpretable feature set for downstream deep learning **(Fig. 1a).** We predicted pathway activities, represented by gene set enrichment scores (GES), from individual tumor genomic profiles and evaluated how well these genomics-derived scores mirrored established expression-based GES from the same samples. **(Fig. 1b).** These pathway activities were then mapped onto hallmark cancer phenotypes, each defined by a collection of related molecular pathways, providing a biologically coherent representation of each tumor **(Fig. 1c).** This representation, the somatic PhenoMap, served as the input feature set for deep learning models, including cancer drug response prediction and prognostic models **(Fig. 1d),** which enables interpretation across multiple layers of biological organization **(Fig. 1e).**

**Figure 1.**
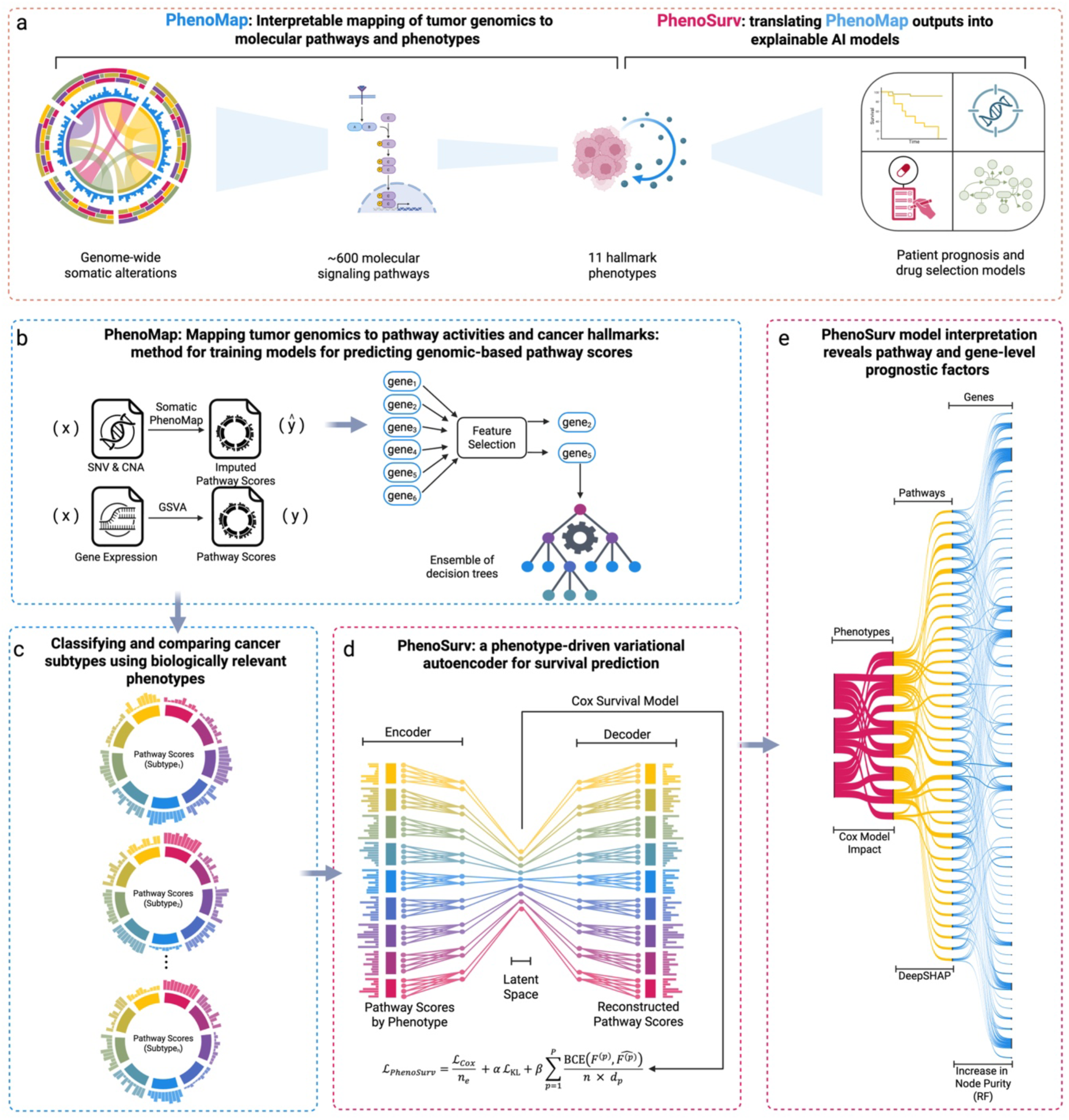
Overview of the PhenoMap framework. **a** PhenoMap integrates molecular pathway activity scores and hallmark cancer phenotypes inferred from somatic alterations. These embeddings can be used in explainable deep learning models, including the PhenoSurv variational autoencoder–based survival model. **b** Data inputs for training PhenoMap pathway models, consisting of paired WES and RNA-seq profiles from TCGA pan-cancer tumors. GES scores for individual tumors were calculated using GSVA and served as the outcome predicted from mutation-based input features. Feature selection and model construction for pathway prediction, using LASSO to identify informative genomic features (SNVs, CNAs) followed by random forest modelling to predict expression-based GES. **c** Representative PhenoMaps illustrating the relative contributions of cancer phenotypes across tumor types. **d** Variational autoencoder architecture underlying PhenoSurv, which leverages PhenoMap-derived pathway and phenotype embeddings to predict survival outcomes. **e** Sankey plot demonstrating model interpretability, showing multi-level feature contributions at the phenotype, pathway, and gene levels.

We evaluated whether SNVs and CNAs could accurately predict expression-derived GES. Using 5-fold cross-validation, the model achieved a mean Pearsons correlation r = 0.51, with the top decile (∼1,000 pathways) reaching r = 0.69 **(Fig. 2a, Suppl. Table 1, Suppl. Fig. 2)**. High performing models (r > 0.75) included cancer-related pathways like cell signaling and immune pathways, while poorly performing models (r < 0.25) were mostly non-cancer pathways **(Suppl. Fig. 3),** indicating pathway-level phenotypic scores can be accurately imputed using tumor mutation data. Nearly 23,000 genes contributed to at least one model (avg. ∼280 genes per pathway) **(Suppl. Fig. 4)**. Of 753 COSMIC cancer census genes, 737 appeared in at least one model, though most contributors (96.9%) were non-COSMIC, suggesting non-canonical genes influence pathway activity **(Suppl. Fig. 5).** The top non-COSMIC genes contributing to multiple pathways included *TTN* (3,112 pathways), *SOX9* (1,651), *DNAH5* (1,480), *SYNE1* (1,052), and *BRINP3* (955), each with well-established roles in their respective phenotypes. *TTN* predicted 98.3% of the immune pathways, consistent with its link to high mutation burden^25^ and response to immunotherapy^26,27^. *SOX9* was prominent in cell death resistance (76.9%) and plasticity (75.0%) and is known for roles in apoptosis and epithelial-to-mesenchymal transition^28,29^. *DNAH5*^30^ and *SYNE1*^31^ both contributed heavily to over 90% of immune pathways. *BRINP3*, associated with proliferation and metastasis^32^, appeared in nearly half of related pathways. Shared genes across top pathways included several COSMIC cancer genes as well as non-canonical genes including, *APC, CASP8, DNAH5, FAT4, IGVL3-1, KMT2D*, MUC16, *MYC, PIK3CA*, and *SOX9* **(Suppl. Fig. 6).**

**Figure 2.**
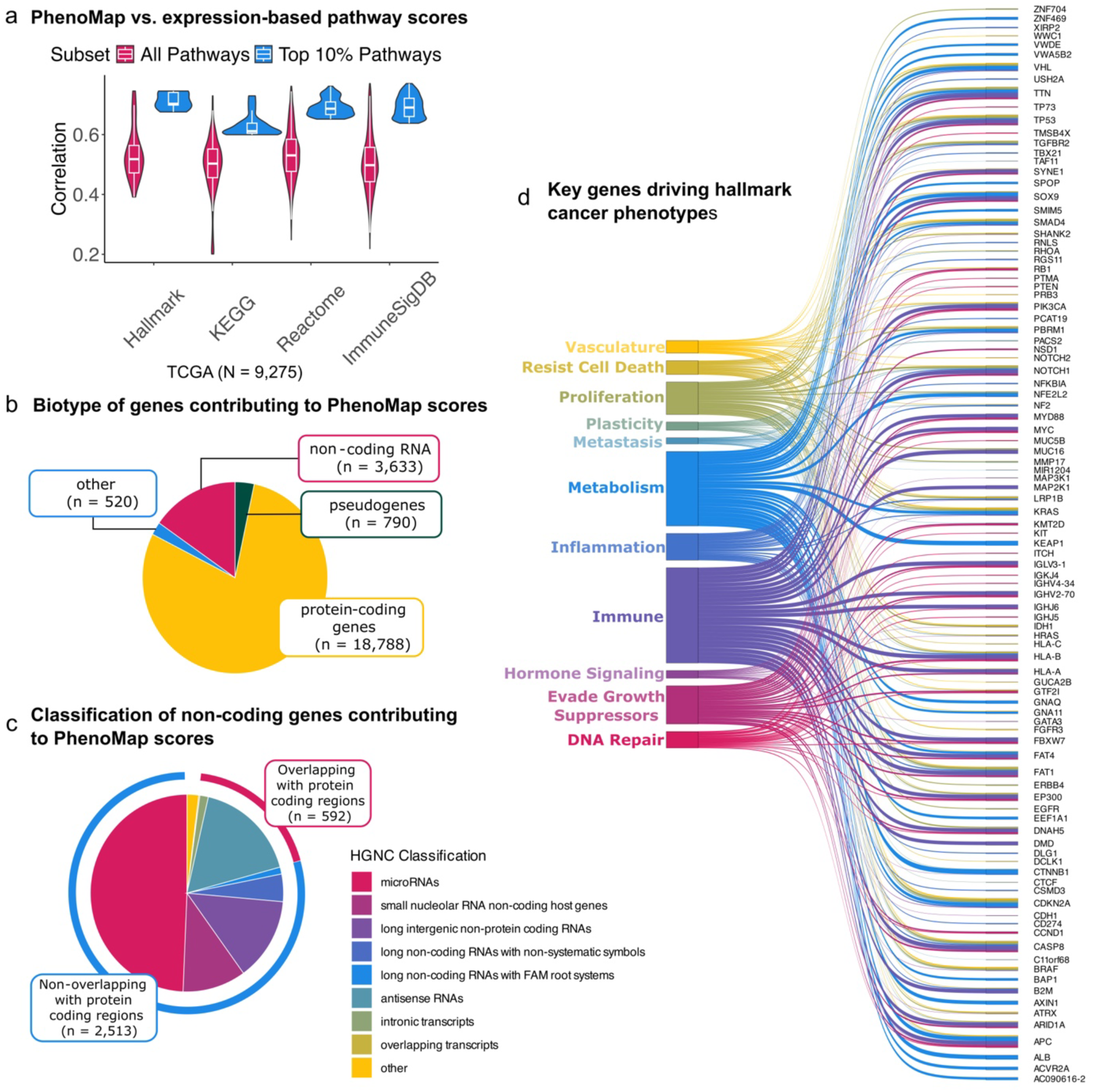
Prediction of pathway scores with genomic data. **a** Violin plots showing distribution of correlations of RNA-based and genomic-based single sample enrichment scores for all pathways (red) and top 10% pathways (blue) with highest correlations from genomic models. **b** Proportion of gene classes selected by LASSO models. **c** Pie chart showing proportion of non-coding RNA based on HGNC nomenclature. **d** Sankey plots showing pathway phenotypes with connections to top 25 genes associated with each phenotype.

Most contributing genes were protein-coding (79.2%), followed by non-coding RNAs (15.3%), pseudogenes (3.3%), and others (2.2%) **(Fig. 2b)**. Top protein-coding genes included *VHL* (3,413 pathways), *KRAS* (3,359), and *TTN* (3,112); non-coding RNAs included *HCCAT5* (724), *PCAT19* (635), and *TUSC7* (448). Notably, most non-coding RNAs (79.2%) did not overlap protein-coding loci, suggesting key roles for long intergenic RNAs in pathway phenotypes **(Fig. 2c)**. We grouped the pathways into 11 phenotypes representing major cancer hallmarks, including DNA repair, evasion of growth suppression, hormone signaling, immune regulation, inflammation, metabolism, metastasis, plasticity, proliferation, resistance to cell death, and vasculature **(Suppl. Table 2).** Shared genes across multiple phenotypes included *TTN, TP53, CASP8, FAT1, APC, PIK3CA, KRAS,* and *SOX9*, while metabolic pathways were distinct, enriched for *KEAP1, CTNNB1,* and *AXIN1*. Immunoglobulin genes appeared across immune, DNA repair, and growth suppression phenotypes, highlighting cross-functional roles **(Fig. 2d).** Our results show that somatic genomic alterations reliably estimated pathway activity with strong concordance to transcriptome-based methods, enabling an interpretable feature layer without expression data. While many contributors were known cancer drivers, the presence of non-coding RNAs, pseudogenes, and cross-phenotype genes highlights a broader functional landscape beyond canonical cancer gene lists.

### PhenoMap accurately classifies cancer subtypes using biologically relevant phenotypes

We assessed PhenoMap’s utility for predictive modeling by generating genomic pathway scores in independent lung, breast, and brain cancer datasets. Models trained on TCGA samples (981 breast, 967 lung, 505 brain) were validated on CPTAC cohorts (121 breast, 109 lung, 95 brain). UMAP projections of pathway scores revealed strong alignment between training and external datasets, with distinct clustering by cancer type. Notably, brain tumors formed a cluster distinct from the two epithelial cancers **(Suppl. Fig. 7a)**. To assess performance, we measured predictive performance with randomized pathway scores (mean r =-0.0018). Models exceeding this baseline included 3,382 for lung, 3,772 for brain, and 4,188 for breast cancer. Of these, models which achieved validation r > 0.25 included 722 for lung, 195 for brain, and 979 for breast cancer **(Suppl. Fig. 7b)**. Principal component analysis (PCA) of the filtered pathway scores revealed clear stratification by subtype within each cancer type. The first two components explained 47.8–54.9% and 7.9–10.5% of total variance, respectively **(Fig. 3a-c).** Pathway-based scores effectively captured intra-cancer heterogeneity, distinguishing clinical subtypes such as basal, HER2+, and luminal A/B in breast cancer; squamous versus adenocarcinoma in lung cancer; and IDH mutation status in brain cancer **(Fig. 1c**; **Fig. 3a-c).** PCA loadings revealed recurrent signals, including DNA unwinding and G2/M checkpoint pathways shared across all three cancer types. Breast cancer was further characterized by cell cycle and DNA replication pathways, reflecting the clinical relevance of CDK4/6 inhibitors^33^ and the contribution of DNA repair genes such as *BRCA*1/2^34^ in breast cancer **(Fig. 3d).** Lung cancer exhibited enrichment for cell cycle and SUMOylation-related pathways in PC1, while PC2 was driven by interleukin and inflammasome signaling, consistent with the substantial role of inflammation as a known mechanism of lung cancer progression^35,36^ **(Fig. 3e)**. In brain cancer, top contributors included apoptotic and inflammatory pathways, two well-known hallmarks of brain cancers^37,38^ **(Figure 3f)**. Together, these results demonstrate that genomic-based pathway models can preserve biologically meaningful structure and enable stratification across diverse cancer contexts.

**Figure 3.**
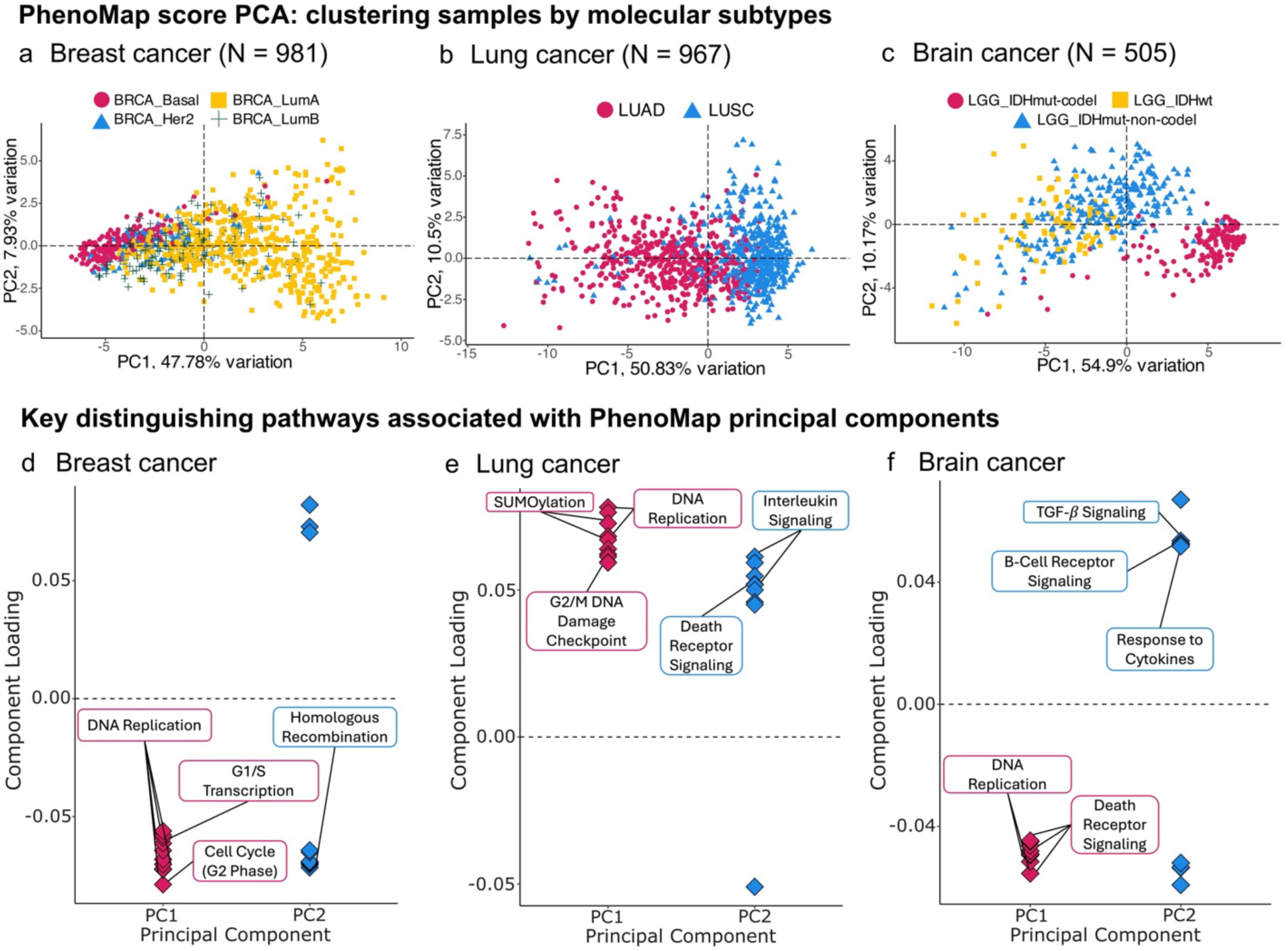
Characterization of cancer subtypes using PhenoMap scores. PCA projections of individual cancer samples in TCGA dataset by subtype in **a** breast (Basal n = 169; Her2 n = 75; LumA n = 496; LumB n = 193), **b** lung (LUAD n = 503; LUSC n = 464), and **c** brain cancers (IDHmut-codel n = 167; IDHmut-noncodel n = 247; IDHwt n = 91). Principal component loadings of top pathways in **d** breast, **e** lung, and **f** brain cancers. Key phenotypes associated with disease subtypes or known resistance mechanisms are highlighted.

### PhenoMap captures tumor heterogeneity and actionable phenotypes

Developing survival and therapy response models in precision oncology requires biologically informative features. To demonstrate clinical relevance, we characterized well-known resistance mechanisms and targetable phenotypes in breast (PI3K/AKT/mTOR, CDK4/6, HER2/EGFR) and lung cancers (HER2/EGFR/ERBB, PI3K/AKT/mTOR, KRAS). PhenoMap pathway scores were significantly correlated with transcriptomic scores across all three resistance mechanisms in breast cancer **(Fig. 4a, Suppl. Fig. 8a).** K-means clustering using transcriptomic or PhenoMap scores grouped cases by dominant signaling pathways, though pathway-driving mutations were distributed across clusters **(Fig. 4b, Suppl. Fig. 8b-d).** PI3K pathway scores showed similar distributions regardless of PI3K mutation status **(Fig. 4c).** Comparing pathways with high principal components contributions revealed three distinct groups with significantly elevated scores for one mechanism using both transcriptomic and PhenoMap scores **(Suppl. Fig. 8e)**. Notably, clustering patients based on PI3K pathway scores provided stronger prognostic value (*p* = 0.0002) than *PIK3CA* mutation status (*p* = 0.72), highlighting limitations of single-gene somatic biomarkers in capturing pathway complexity **(Fig. 4d).** Cluster assignments across breast cancer subtypes aligned with known actionable phenotypes. For instance, luminal A tumors were enriched for PI3K pathway, while HER2-positive tumors were enriched for EGFR pathway, and highly proliferative basal tumors were enriched for CDK4/6 signaling **(Fig. 4e).**

**Figure 4.**
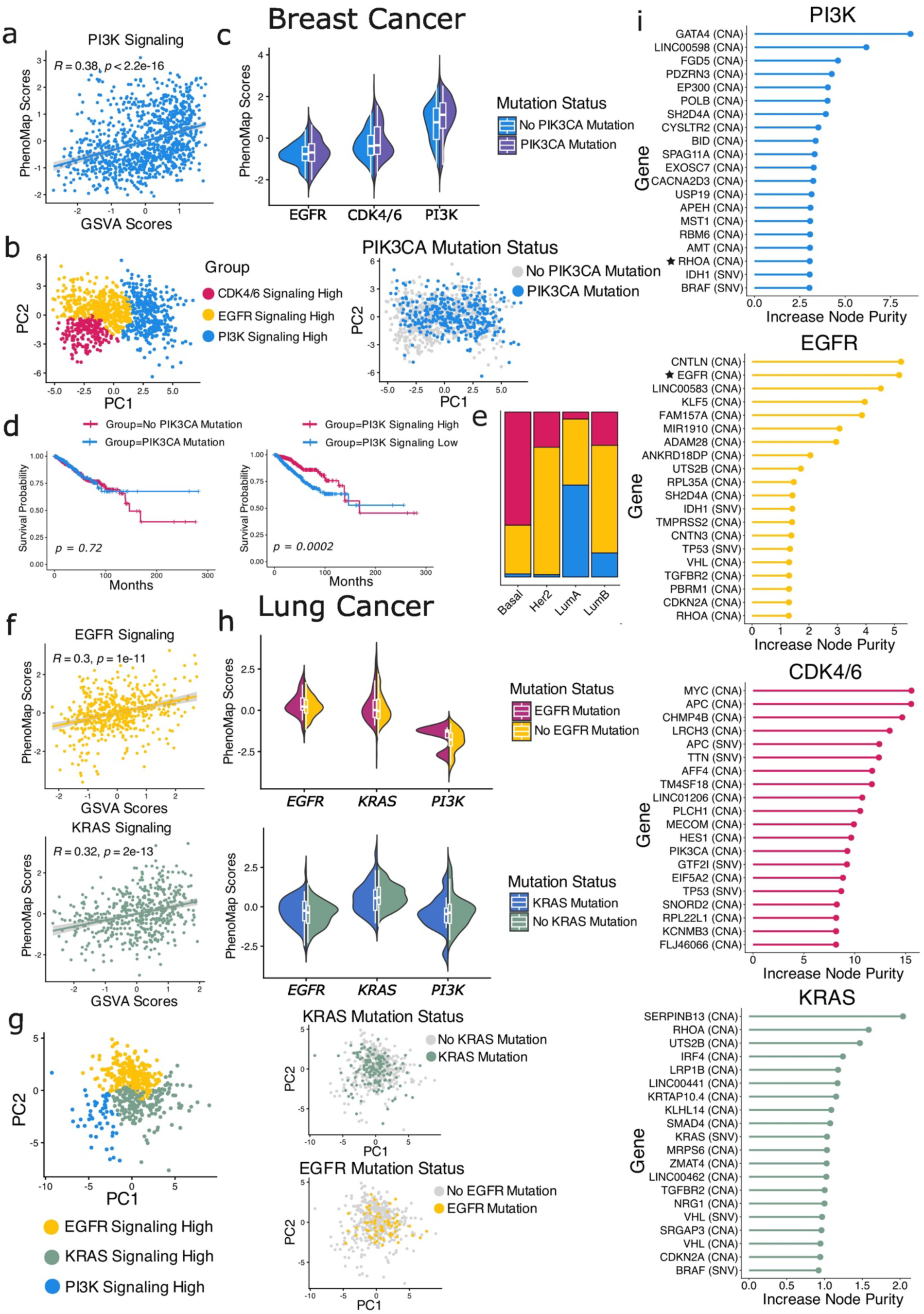
Capturing targetable phenotypes in breast and lung cancer. **a** Spearman correlation of GSVA GES and PhenoMap pathway scores in breast cancer. **b** PCA projections of breast cancer clusters based on most activated signaling pathway. Clusters from PhenoMap scores were EGFR signaling high (n = 206), CDK4/6 signaling high (n = 197), and PI3K signaling high (n = 331). **c** Split violin plot of PhenoMap PI3K Signaling scores by cluster in patients with or without *PI3KCA* mutation. **d** Progression-free survival in breast cancer cases with or without *PIK3CA* mutation (left) or in breast cancer cases with PhenoMap scores above or below mean value in the Reactome PI3K/AKT activation pathway (right). **e** Stacked bar chart of proportions of patients per breast cancer subtype in each cluster. **f** Spearman correlation of GSVA and PhenoMap pathway scores in lung cancer. **g** PCA projections of lung adenocarcinoma clusters based on most activated signaling pathway. Clusters from PhenoMap scores were EGFR signaling high (n = 255), KRAS signaling high (n = 196), and PI3K signaling high (n = 52). **h** Split violin plot of PhenoMap KRAS and EGFR Signaling scores by cluster in patients with or without relevant mutation. **i** Top 20 most important features in PhenoMap score calculation for each pathway. Stars indicate genes also included in GSVA method.

Strong concordance of PhenoMap vs. expression-based GES was also observed across all three resistance mechanisms in lung cancer **(Fig. 4f, Supplementary Figure 9a).** Clustering of lung cancer cases based on resistance mechanisms and targetable phenotypes also revealed distinction between phenotypic states **(Fig. 4g, Suppl. Fig. 9b,d,e).** Additionally, comparison of representative pathways for each mechanism showed significantly higher scores compared to alternate mechanisms **(Suppl. Fig. 9c).** EGFR, CDK4/6, and PI3K pathway scores showed similar distributions within each cluster regardless of related gene mutation status **(Fig. 4h),** highlighting that single-gene biomarkers may be insufficient for assessment of patient prognosis.

Top 20 features per pathway model included several high-importance genes not traditionally linked to these resistance mechanisms **(Fig. 4i).** *RHOA* contributed to PI3K signaling for both PhenoMap and GSVA, alongside *EGFR* in the EGFR pathway; the remaining featured genes in PhenoMap were absent from the corresponding MSigDB signatures used in expression-based GES calculations. G1/S transition featured *MYC, APC, CHMP4B*, and *AFF4*, genes with established roles in cell cycle regulation^39–42^. EGFR signaling included *CNTLN* – a gene linked to cancer progression^43^ – followed by *EGFR, KLF5*, and multiple long-coding and microRNAs (*LINC00583, FAM157A*, and *MIR1910*). PI3K/AKT activation highlighted genes with emerging evidence of involvement, such as *GATA4*^44^, *LINC00598*^45^, *FGD5*^46^, and *PDZRN3*^47^. KRAS signaling incorporated known players like *RHOA*^48^, *LRP1B*^49^, and *SMAD4*^50^, as well as understudied genes including *SERPINB13, UTS2B*, and *IRF4*. PhenoMap-derived pathway scores capture biologically relevant resistance mechanisms, offering richer prognostic context than single-gene mutations and identifying somatic contributors beyond canonical pathway genes.

### PhenoSurv: a phenotype-driven VAE reveals NOTCH1 signaling and *SMARCA4* as key breast cancer prognostic factors

Recent AI/ML advances enable generative models for clinical outcome prediction, but poor interpretability limits their clinical use. To address this, we developed PhenoSurv, a variational autoencoder that embeds phenotype-grouped pathway scores from PhenoMap into latent dimensions optimized with phenotype reconstruction, KL divergence, and Cox loss. Unlike prior VAEs, PhenoSurv integrates biological information to produce interpretable latent features highlighting multi-phenotypic vulnerabilities. **(Fig. 1d–e).** Moreover, clinical genomic sequencing data typically includes a limited set of genes, which requires careful evaluation of the predictive capability of smaller subset of somatic variants. To address this, we evaluated PhenoSurv on MSK IMPACT, which includes ∼400 genes.

PhenoSurv models were trained on TCGA and MSK cohorts to predict disease-free or progression-free survival (DFS/PFS) for lung adenocarcinoma (TCGA n=246; MSK n=584), brain cancer (TCGA n=504; MSK n=434), and breast cancer subtypes: HR+/HER2– (TCGA n=398; MSK n=1,049), HER2+ (TCGA n=143; MSK n=146), and TNBC (TCGA n=101; MSK n=123).

PhenoSurv outperformed random survival forests and DeepSurv trained on raw genomic data, achieving mean concordance indices of 0.7349 (TCGA) and 0.5434 (MSK) versus 0.4112–0.5131 (TCGA) and 0.4897–0.5009 (MSK). PhenoSurv achieved significantly higher concordance indices compared to random survival forest (*p* = 0.0048) and DeepSurv (*p* = 0.00006). **(Suppl. Fig. 10a, Suppl. Table 3).** Although random survival forests also provide interpretability via feature importance, PhenoSurv demonstrated substantially more consistent identification of survival-relevant features. Across the five trained models for each method, the mean Spearman correlation was higher for PhenoSurv (TCGA: 0.5846; MSK: 0.3578) compared with random survival forests (TCGA: 0.0960; MSK: 0.3051), indicating greater reproducibility and more reliable identification of prognostic biomarkers **(Suppl. Fig. 10b)**. Univariate Cox analysis of latent dimensions showed proliferation as the strongest predictor across all subtypes, followed by metabolism and vasculature in HR+/HER2–, DNA repair and metastasis in HER2+, and metabolism and inflammation in TNBC **(Fig. 5a, Suppl. Fig. 10c).**

**Figure 5.**
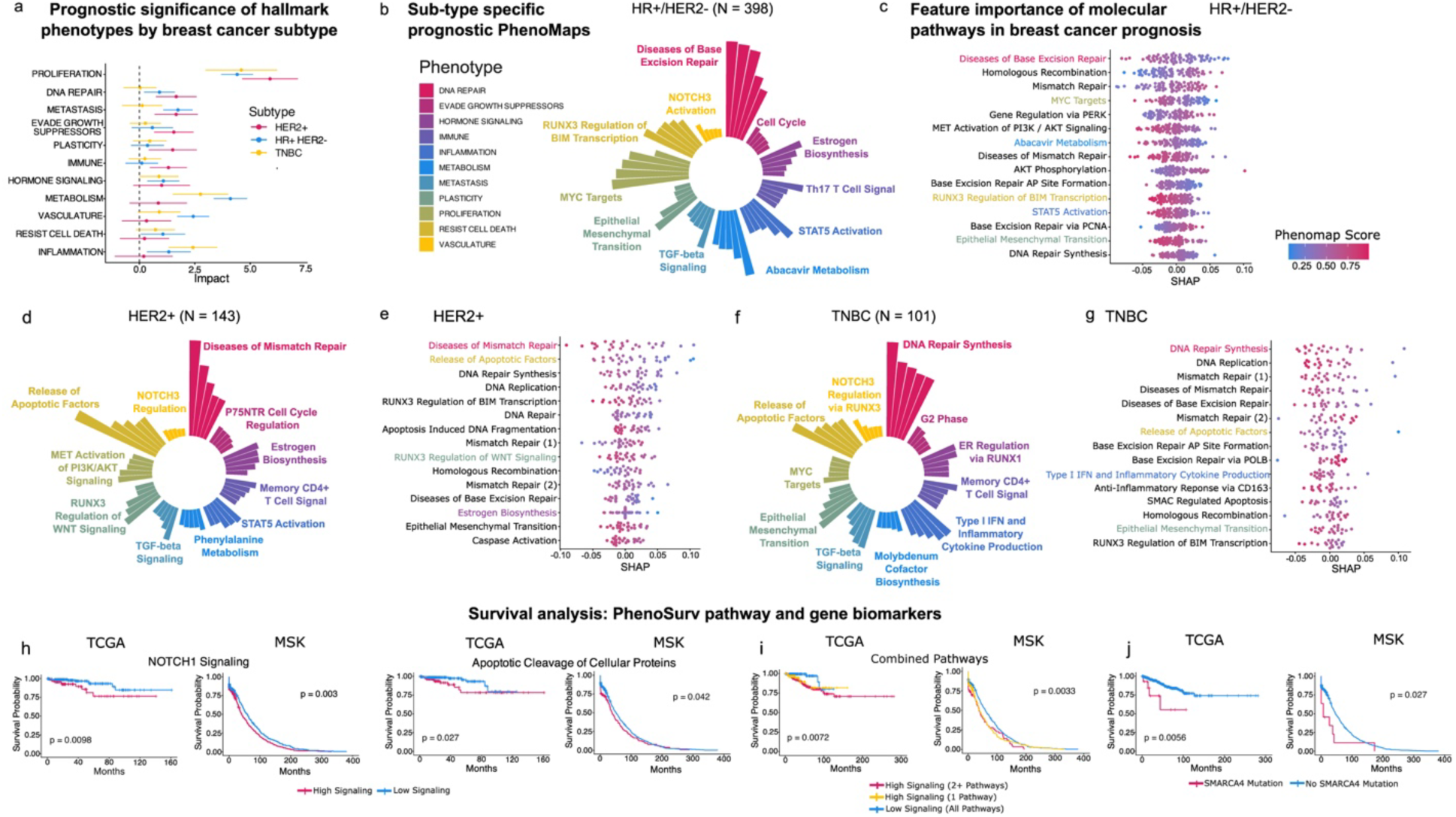
Pathway phenotype-driven deep survival model in breast cancer by subtypea. Comparison of hazard ratios of phenotype-based latent dimensions from trained PhenoSurv models. Mean SHAP values of the top 5 pathways from each phenotype in **b** HR+/HER2-, **d** HER2+, and **f** Triple negative breast cancers. SHAP values of top 15 pathways overall in TCGA dataset in **c** HR+/HER2-, **e** HER2+, and **g** Triple negative breast cancers. The top pathways for each phenotype are highlighted by color. **h** Kaplan-Meier disease-free survival plot of patients with high vs low signaling in TCGA and MSK breast cancer datasets. **i** Kaplan-Meier disease-free survival of breast cancer patients with multiple high signaling pathways in TCGA and MSK breast cancer datasets. **j** Kaplan-Meier disease-free survival plot of patients with or without SMARCA4 mutation in TCGA and MSK breast cancer datasets.

To interpret survival predictions at the pathway level, we computed SHAP values across subtype-specific models. Common high-impact pathways included DNA repair (base excision, mismatch repair, homologous recombination), epithelial–mesenchymal transition, and apoptosis via *RUNX3* regulation of BIM transcription. HR+/HER2– tumors showed unique enrichment for Abacavir metabolism, MYC targets, and PI3K/AKT activation. Among inflammation pathways, *STAT5* activation ranked in the top five for all subtypes, being most prominent in HR+/HER2– and HER2+ cancers. In TNBC, Type I IFN signaling and inflammatory cytokine production dominated, followed by CD163-induced anti-inflammatory response **(Fig. 5b-g, Suppl. Fig. 10d).**

PhenoSurv’s interpretability enables identification of multi-level biomarkers from trained survival models. Screening the top ten pathways from each phenotype for HR+/HER2- breast cancer revealed two significant pathway-level biomarkers: NOTCH1 signaling (TCGA: *p* = 0.0098; MSK: *p* = 0.003) and Apoptotic cleavage of cellular proteins (TCGA: *p* = 0.027; MSK: *p* = 0.042), both associated with distinct disease-free survival cohorts **(Fig. 5h)**. While deregulation of NOTCH1 signaling and apoptosis are well-documented in breast cancer biology^51,52^, clinically actionable markers from these pathways remain limited. There was a significant difference in disease-free survival across patients with low signaling in both the *NOTCH1* and apoptotic pathways, patients with high signaling in one pathway, and those with high signaling in both pathways (TCGA: *p* = 0.0072; MSK: *p* = 0.0033) **(Fig. 5i)**. Our model shows that combining these pathways further amplified survival differences between patients with low versus high signaling, highlighting their importance in HR+ breast cancer patient prognosis. Gene-level screening of PhenoMap components identified *SMARCA4*, a contributor to apoptotic cleavage of cellular proteins, as significantly associated with worse survival when mutated (TCGA: *p* = 0.0056; MSK: *p* = 0.027) **(Fig. 5j)**. Although not an established biomarker for HR+ breast cancer, *SMARCA4* is an important part of the SWI/SNF chromatin remodeling complex, and its loss has been linked to aggressive cancer phenotypes^53^. Moreover, SWI/SNF complex controls the transcriptional activity of NOTCH1 signaling through regulation of enhancer accessibility^54,55^. Our findings again highlight PhenoMap’s translational potential through discovery of biologically relevant multi-level predictors of patient prognosis.

### PhenoSurv identifies TGF-β/SMAD5 activation as a key prognostic pathway in lung adenocarcinoma

We next evaluated PhenoSurv on lung adenocarcinoma datasets for disease-free and recurrence-free survival in TCGA and MSK, respectively. In lung adenocarcinoma, PhenoSurv outperformed conventional models trained on raw genomic data, achieving a mean CI of 0.5500 (TCGA) and 0.6238 (MSK), compared to CIs of 0.4730–0.5056 (TCGA) and 0.4946-0.54726 (MSK) from random survival forests and DeepSurv models **(Suppl. Fig. 11a, Suppl. Table 3).** PhenoSurv again showed superior reproducibility of ranked survival-relevant features across models. Spearman correlation was higher for PhenoSurv (TCGA: 0.5138; MSK: 0.5178) compared with random survival forests (TCGA: 0.0686; MSK: 0.3864) **(Suppl. Fig. 11b)**. The most influential phenotypes included vasculature, metabolism, and metastasis **(Fig. 6a, Suppl. Fig. 11c).**

**Figure 6.**
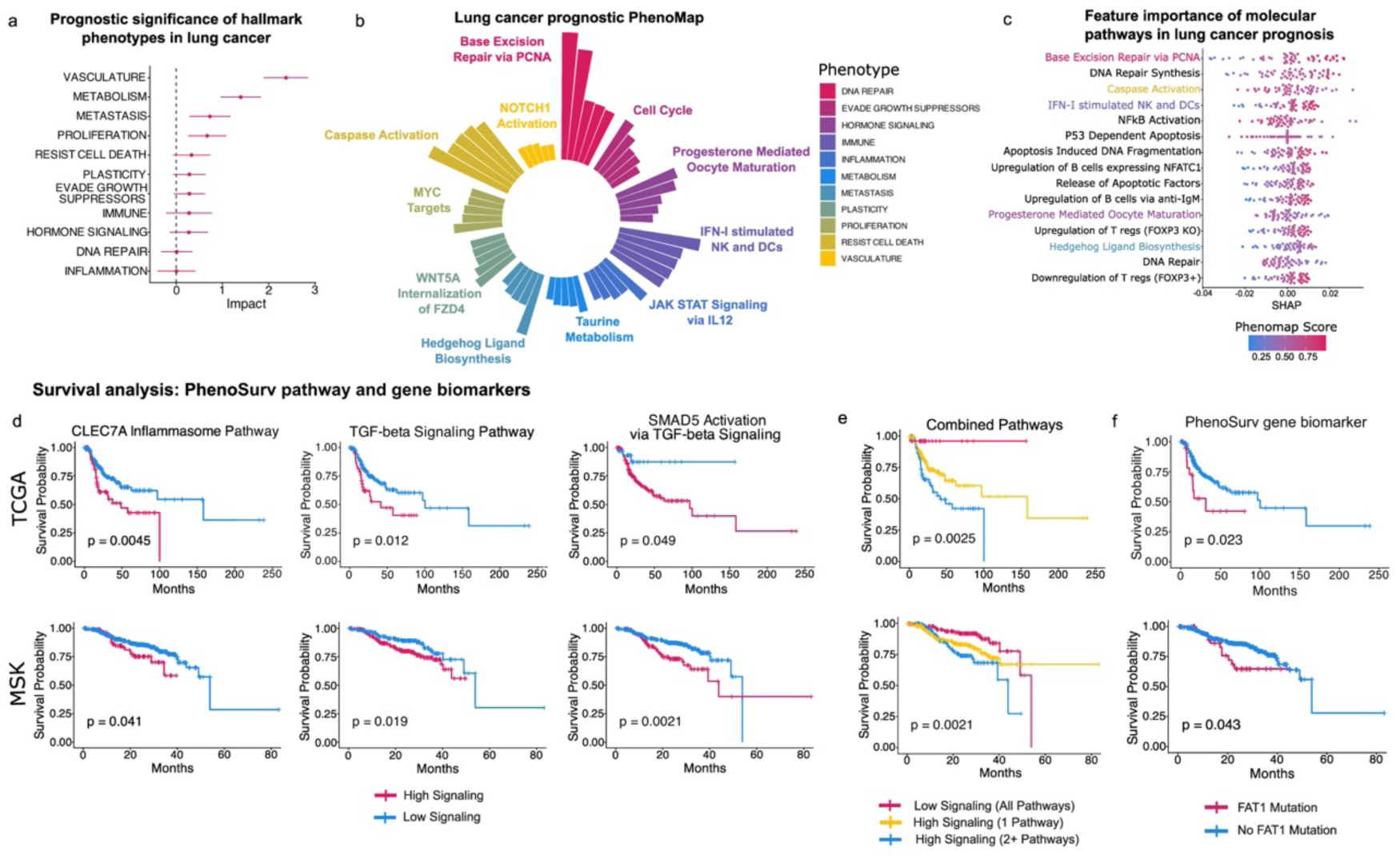
Pathway phenotype-driven deep survival model in lung cancer. (TCGA n = 246, MSK n = 434). **a** Comparison of hazard ratios of phenotype-based latent dimensions from trained PhenoSurv models. **b** Mean SHAP values of top 5 pathways from each phenotype. **c** Beeswarm plot of SHAP values of top 15 pathways overall in TCGA dataset. The top pathways for each phenotype are highlighted by color. **d** Kaplan-Meier survival plot of patients with high vs low signaling in TCGA and MSK lung cancer datasets. **e** Kaplan-Meier survival plot of patients based on combination of three pathway markers. **F** Kaplan-Meier survival curves of patients with or without FAT1 mutation in TCGA and MSK lung cancer datasets.

SHAP analysis of top pathways per phenotype showed DNA repair pathways were relevant in lung cancer but less influential than in breast cancer. Immune-related pathways were more prominent in lung cancer, consistent with its immunologically active profile^56^. Key immune signatures included IFN-I stimulated NK and dendritic cells, B-cell anti-IgM response, and multiple T-cell gene sets upregulated with FOXP3 knockout—all associated with worse prognosis **(Fig. 6b-c, Suppl. Fig. 11d).** Notably, prior studies link higher FOXP3+ T-cell levels to improved immunotherapy response in lung cancer patients^57^.

Three pathway biomarkers for lung adenocarcinoma stratified patients by disease-free survival (TCGA) and recurrence-free survival (MSK): *CLEC7A* inflammasome (TCGA: *p* = 0.0045; MSK: *p* = 0.041), TGF-β signaling (TCGA: *p* = 0.012; MSK: *p* = 0.019), and *SMAD5* activation via TGF-β (TCGA: *p* = 0.049; MSK: *p* = 0.0021) **(Fig. 6d)**. Combining these biomarkers revealed a clear survival gradient: patients with all favorable markers had the best outcomes, those with one unfavorable marker had intermediate survival, and patients with multiple unfavorable markers had the poorest outcomes (TCGA: *p* = 0.0025; MSK: *p* = 0.0021) **(Fig. 6e).** Both TGF-β signaling and *SMAD5* activation are linked with lung adenocarcinoma metastasis^58,59^ and are being explored as therapeutic targets. Although the role of *CLEC7A* in triggering inflammatory response is well-characterized, its precise role is not well-established in lung cancer. Gene-level analysis highlighted the contributions of *FAT1* gene in the *CLEC7A* and TGF-β pathways, and its mutations were correlated with worse survival (TCGA: *p* = 0.023; MSK: *p* = 0.043) **(Fig. 6f).** *FAT1* has a potential role in activating TGF- β signaling, while cell adhesion molecules also regulate inflammation^60^. Although *FAT1* is not an established clinical biomarker, its diverse roles in tumor progression and emerging evidence as an immunotherapy marker highlight its potential importance in lung cancer prognosis^61,62^.

### PhenoSurv identifies multiple targetable phenotypes as prognostic factors in glioblastoma

In brain cancer, PhenoSurv achieved mean C-indices of 0.7042 (TCGA) and 0.7573 (MSK), compared to benchmarks (TCGA: 0.6304–0.691; MSK: 0.6047–0.7205). PhenoSurv achieved significantly higher concordance indices compared to DeepSurv (*p* = 0.0253). **(Suppl. Fig. 12a, Suppl. Table 3)**. Mean spearman correlation was also higher for PhenoSurv (TCGA: 0.7807; MSK: 0.3473) compared with random survival forests (TCGA: 0.3765; MSK: 0.0365) for the brain cancer models **(Suppl. Fig. 12b)**. Top phenotypes included vasculature, proliferation, and evasion of growth suppressors **(Fig. 7a, Suppl Fig. 12c)**, with the G2/M checkpoint showing particularly high relevance. Consistent with other cancer types, DNA repair and apoptotic pathways were prominent, while WNT signaling, a key brain cancer pathway, contributed more strongly than in other cancers. Notably, *IRAK4* deficiency ranked among the top contributors, supportive of ongoing investigations of IRAK4 inhibitors for brain metastases^63^ **(Fig. 7b-c, Suppl. Fig. 12d).**

**Figure 7.**
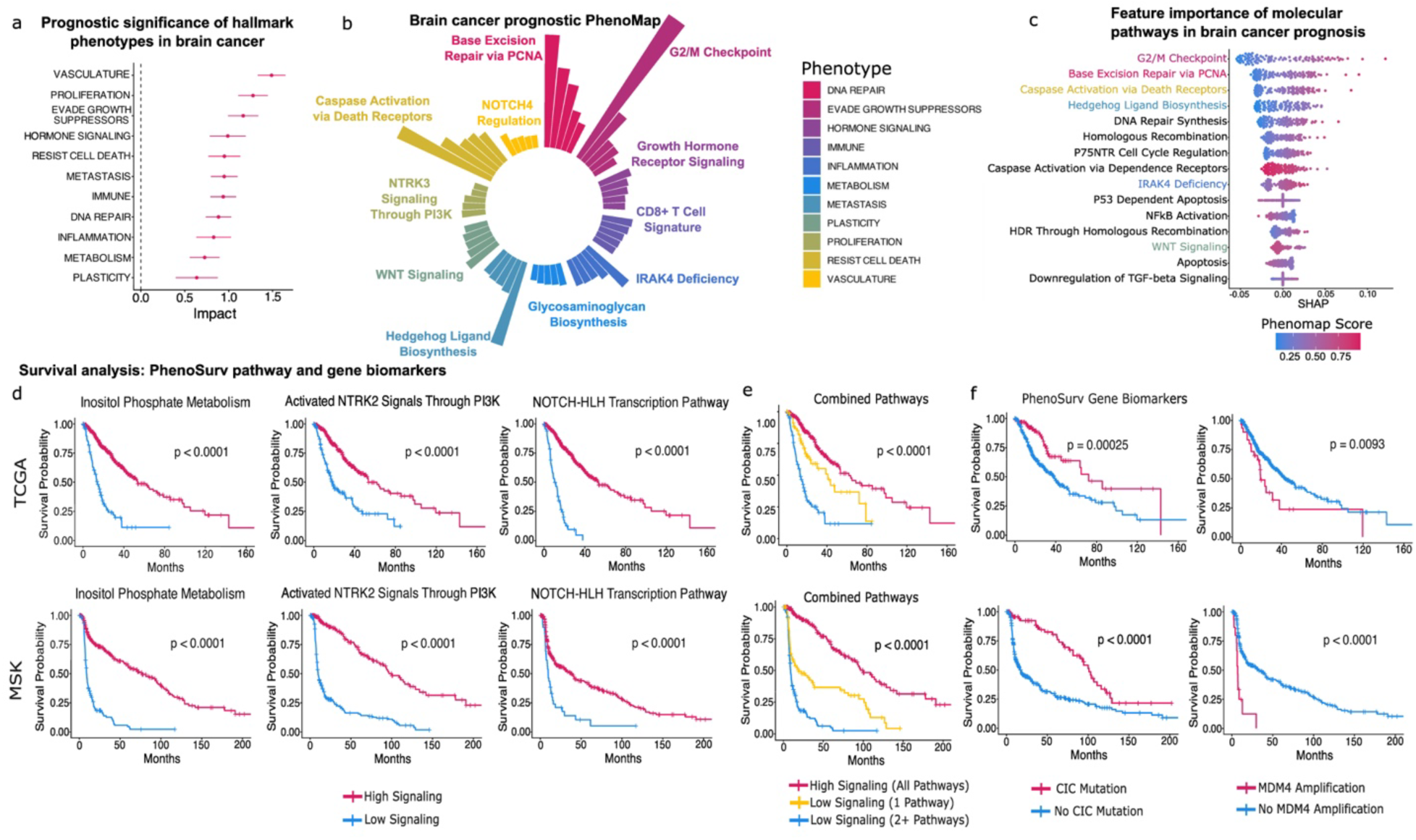
Pathway phenotype-driven deep survival model in brain cancer. (TCGA n = 504, MSK n = 343). **a** Comparison of hazard ratios of phenotype-based latent dimensions from trained PhenoSurv models. **b** Mean SHAP values of top 5 pathways from each phenotype. **c** Beeswarm plot of SHAP values of top 15 pathways overall in TCGA dataset. The top pathways for each phenotype are highlighted by color. **d** Kaplan-Meier survival plot of patients with high vs low signaling in TCGA and MSK brain cancer datasets. **e** Kaplan-Meier survival plot of patients based on combination of three pathway markers. **f** Kaplan-Meier survival plot of patients with or without *CIC* mutation or *MDM4* amplification in TCGA and MSK brain cancer datasets.

Over half of the top pathways screened for brain cancer (58 total) were significant biomarkers, stratifying patients by progression-free survival in both TCGA and MSK datasets **(Suppl. Table 4).** The three strongest pathways based on average chi-squared values were inositol phosphate metabolism, activated *NTRK2* signaling via PI3K, and NOTCH-HLH transcription (all *p* < 0.0001 for TCGA and MSK) **(Fig. 7d)**. Notably, inositol treatment is under investigation for brain cancer^64^, and PI3K and NOTCH signaling are well-established drivers^65,66^. Consistent with other cancer types, survival decreased progressively with the number of poor prognostic markers **(Fig. 7e).** Eight gene-level biomarkers were identified, including known markers *IDH1/2, EGFR*, and *PTEN* **(Suppl. Fig. 12e).** The role of *CIC* in brain cancer is less studied compared to other markers, though it has been linked to MAPK activation and may represent a therapeutic target^67,68^. Notably, *CIC* mutations were associated with improved survival (TCGA: *p* = 0.00025; MSK: *p* < 0.0001). *MDM4* amplifications were associated with worse survival (TCGA: *p* = 0.0093; MSK: *p* < 0.0001), consistent with its role in the TP53/MDM2/MDM4/p14AR pathway and emerging evidence for *MDM2* as a prognostic biomarker^69^ **(Fig. 7f).**

Together, these results demonstrate that PhenoSurv not only achieves accurate survival predictions but also uniquely enables multiscale biological interpretability—from phenotype to pathway to gene. This interpretive capacity provides mechanistic insight into cancer progression and highlights clinically relevant genomic features often missed by black-box models. PhenoSurv provides a powerful framework for precision oncology, enabling both prognostic modeling and discovery of actionable biomarkers.

## DISCUSSION

In this study, we present a novel framework for developing biologically interpretable prognostic models using somatic genomic data. PhenoMap accurately predicts hundreds of pathway activity scores from mutation and copy number data, showing high concordance with RNA-based GSVA scores across test and validation cohorts. While widely used tools such as Integrative Genomics Browser (IGV) and the Catalogue for Somatic Mutations in Cancer (COSMIC) provide visualization and mutation signature insights, they offer limited biological interpretability. In contrast, expression-based tools like GSVA, despite their popularity, lack genomic equivalents. This gap is critical, as RNA sequencing remains less accessible in both clinical and research settings compared to genomic profiling. Our approach addresses this need by enabling pathway-level interpretation directly from genomic data.

PhenoMap revealed that a broad spectrum of genes, including those not typically classified as oncogenic, such as noncoding RNAs and pseudogenes, contribute meaningfully to pathway-level predictions. This highlights the biological relevance of many genes often overlooked in clinical genomic analyses. Genes appearing across hundreds of pathway models likely reflect true biological significance; for example, HLA and immunoglobulin genes ranked highly in immune and DNA repair pathways, consistent with evidence linking DNA damage to antigen presentation^70^. Together, these findings emphasize PhenoMap’s utility not only in interpreting somatic genomic data but also in uncovering novel gene-pathway associations relevant to cancer biology.

PhenoMap introduces a powerful approach for analyzing somatic genomic data by generating biologically interpretable pathway activity scores. These scores provide a foundation for downstream analyses, including patient stratification and tumor-type comparisons. Dimensionality reduction using UMAP and PCA demonstrated that PhenoMap-derived scores effectively separate cancer types, subtypes, and resistance mechanisms, revealing molecular patterns and subgroup-specific biology across diverse cohorts. Current clinical practice relies heavily on single-gene biomarkers for guiding targeted therapies; however, these markers capture only a fraction of the biological context and often fail to predict treatment response. For example, although *PIK3CA* mutations were common in the TCGA breast cancer cohort, many patients harboring these mutations clustered within groups exhibiting high EGFR or CDK4/6 signaling scores—despite lacking mutations in those pathways. This suggests that treatment decisions based solely on single-gene markers may lead to suboptimal regimens, emphasizing the need for pathway-level insights to improve precision oncology.

Building on PhenoMap’s ability to capture key cancer-related pathways, we developed PhenoSurv, an interpretable deep learning framework for cancer prognosis. Unlike traditional models, PhenoSurv integrates biologically informed phenotypic features with deep learning architecture to identify multi-phenotype drivers of cancer. Applied to breast, lung, and brain cancers, PhenoSurv consistently outperformed existing survival models. By embedding phenotype-level features before Cox survival modeling, PhenoSurv enables interpretation across phenotypes, pathways, and genes—addressing a major limitation of current black-box approaches. This multiscale interpretability allows PhenoSurv to train accurate prognostic models while uncovering novel biomarkers. In HR+/HER2– breast cancer, pathways such as *NOTCH1* signaling and apoptotic processes stratified patients by outcome, reflecting underlying mechanisms related to estrogen regulation and cell death. Although these pathways and the *SMARCA4* gene are not established clinical biomarkers in breast cancer, existing literature provides strong biological rationale linking them to more aggressive disease phenotypes^51–53^. Importantly, the features identified by PhenoSurv function not only as prognostic biomarkers but also as potential therapeutic targets. Investigational treatments already exist for several of these markers, including nirogacestat—a γ-secretase inhibitor that blocks Notch activation and is currently being evaluated in desmoid tumors^71^. Additionally, early-phase clinical trials have begun testing selective *SMARCA2* inhibitors for tumors harboring *SMARCA4* alterations^72^.

In lung adenocarcinoma, the *CLEC7A* inflammasome pathway and TGF-β signaling have emerged as key prognostic features, consistent with the tumor’s immunologically active nature and resistance to therapy^58,59^. Although *CLEC7A*-targeted approaches are not yet in clinical trials, preclinical studies indicate that activating *CLEC7A* (Dectin-1) can enhance anti-tumor immunity and may help overcome acquired resistance to immune checkpoint blockade in melanoma^73^. The strong prognostic relevance of *CLEC7A* in this cohort suggests that targeting this pathway could be a promising strategy for addressing therapeutic resistance in immunologically “hot” tumors and warrants further investigation. In contrast, TGF-β signaling is a well-established driver of cancer progression, and multiple inhibitors are advancing through clinical development. Several TGF-β inhibitors are expected to reach clinical availability in the coming years, including AdAPT-001, now in Phase II trials (NCT04673942)^74^, and STP707, currently in Phase I (NCT05037149)^75^. The biomarkers identified by PhenoSurv in this study highlight specific patient subgroups who may be strong candidates for these emerging therapeutic agents.

Brain tumors displayed distinct pathway signatures, revealing unique molecular drivers and a substantial set of novel prognostic pathway biomarkers. Inositol metabolism emerged as a prominent biomarker, alongside additional metabolic pathways taurine and hypotaurine metabolism, glycogen metabolism, glycosaminoglycan biosynthesis, steroid hormone metabolism, polyamine metabolism, and peptide hormone biosynthesis. Growing interest in metabolic reprogramming has accelerated the development of therapies that target cancer metabolism and sensitize tumors to standard treatments^64,76^. The variety of significant metabolism biomarkers identified by PhenoSurv demonstrates that the framework can not only uncover new metabolic vulnerabilities but also help guide personalized metabolic therapy. Combining multiple pathway biomarkers further amplified survival stratification, showing the value of integrative risk assessment. Gene-level analysis identified well-established cancer markers, including IDH1/2, EGFR, and PTEN, highlighting PhenoSurv’s ability to recover clinically relevant biomarkers. It also surfaced less frequently used but biologically meaningful candidates, such as CIC and MDM4, both associated with more aggressive tumor phenotypes^67–69^. Collectively, these results illustrate how PhenoSurv integrates predictive performance with mechanistic insight, enabling the discovery of actionable biomarkers that extend beyond traditional single-gene approaches.

Although identifying aggressive disease is critical for guiding clinicians toward informed treatment decisions, a key limitation of this study is the heterogeneity of the cohorts, which include patients across different stages and with varied treatment histories. Future applications of the PhenoMap–PhenoSurv framework will focus on training models using cohorts with standardized treatment regimens or well-defined therapeutic contexts. Such refinement will enable the identification of biomarkers predictive of response to specific drugs and improve outcome modeling for targeted therapies. Ultimately, this approach could support personalized treatment planning by linking phenotypic states to therapy-specific prognostic signatures, thereby advancing precision oncology beyond generalized survival prediction.

## CONCLUSIONS

Together, PhenoMap and PhenoSurv provide an opportunity to enhance the utility of existing clinical genomic tests by mapping alterations to actionable biological mechanisms. PhenoMap enables biologically grounded interpretation of mutation and copy number alterations through pathway-level activity scores and broad cancer phenotypes, overcoming the limited interpretability of traditional genomic tools at the single gene level and the accessibility barriers of RNA-based approaches. By leveraging these pathway and phenotype scores, PhenoSurv extends this interpretability to survival prediction, revealing phenotype-, pathway, and gene-specific contributions to patient outcomes. These tools help establish a path forward for more interpretable, biology-driven machine learning models in oncology and lay the groundwork for more precise stratification of patients. Increasing the integration of somatic genomic data into clinical decision-making may aid the development of targeted therapeutic strategies aligned with individual tumor biology.

## METHODS

### Pan-cancer Training Dataset Curation

Pan-cancer genomic, transcriptomic, and clinical data were downloaded from Genomic Data Commons on 1/13/2025. A total of 9,276 samples were used for training and testing the PhenoMap models including breast (n = 994), lung (n = 969), gastrointestinal (n = 1,579), head and neck (n = 565), bladder (n = 403), gynecological (n = 1,040), brain (n = 655), kidney (n = 691), endocrine/neuroendocrine (n = 758), prostate (n = 651), melanoma (n = 443), testicular (n = 127), sarcoma (n = 230), and lymphoma (n = 37) cancer types. Cancer types of a small subset of samples (n = 134) were not found within the clinical data. The GDC processing pipeline for whole genome sequencing aligned raw FASTQ files with BWA-MEM using the human reference genome GRCh38.d1.vd1. The GATK MuTect2 pipeline was used for somatic variant calling. Copy number alterations were called using the GATK4 CNA pipeline. The mRNA analysis pipeline used STAR 2 TwoPass for alignment to GRCh38 reference genome followed by expression transformation using FPKM calculations. Detailed description of all sequencing data processing methods can be found in the GDC data user’s guide^77^. The mutation data file was restructured from VCF format to binary gene-level data. Variants with Ensembl Sequence Ontology terms classified as either high or moderate by IMPACT rating were included in the curated data. For each gene, a value of either 1 or 0 was used to represent whether the patient had one or more impactful mutations on that gene (1), or no impactful mutations on that gene (0). Copy number data was represented by GISTIC gene-level copy number threshold calls^78^.

### Validation Dataset Curation

To validate the correlation of the genomic-based pathways to RNA-based pathway scores, SNV, CNA, and gene expression data from breast (n = 121), lung (n = 109), and brain (n = 95) cancers from the Clinical Proteomic Tumor Analysis Consortium (CPTAC) were downloaded from cBioPortal on 4/8/2025. CPTAC genomic and transcriptomic datasets were pre-processed with the same GDC processing pipeline as used for the pan-cancer cohort. SNV and CNA datasets were converted to gene-level binary mutation and copy number threshold calls, and gene expression data was used to calculate single sample pathway-level scores using the same methods as above.

For replicating our phenotype-driven survival VAE model (PhenoSurv), MSK-IMPACT targeted sequencing datasets from breast (n = 1,318)^79^, lung (n = 584)^80^, and brain (n = 646)^81^ cancers were downloaded from cBioPortal on 4/16/2025. Samples sequenced based on the 410 gene panel or larger gene panels were included in the study. Similarly to the GDC pipeline, the MSK-IMPACT data processing pipeline used BWA for alignment using reference genome GRCh37, and GATK for base quality recalibration. Somatic mutation calling was preformed using MuTect, SomaticIndelDetector (GATK), and PINDEL. Copy number aberrations were found by comparing the coverage of tumor compared to normal samples in targeted regions. Detailed description of MSK-IMPACT sequencing data processing can be found at https://impact-pipeline.readthedocs.io/en/latest/. SNV and CNA datasets were further curated for our analysis using the same methods as the pancancer datasets.

### Expression-based gene set enrichment scores calculation

mRNA data represented by gene-level expression was transformed using FPKM calculations. Expression data was used for gene set variational analysis (GSVA) to calculate single sample pathway-level scores from MSigDB Hallmark, KEGG (Legacy), Reactome, and ImmuneSigDB pathway collections using the R packages *GSVA*^82^ and *msigdbr*. The *gsva* method used for pathway score calculations is a non-parametric method which calculates expression-level statistics to bring gene expression profiles to a common scale. The enrichment scores were calculated as the magnitude difference between the largest positive and negative Kolmogorov-Smirnov random walk deviations^82^.

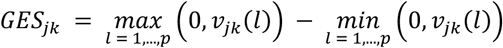

### Genomic-based pathway scoring

To establish the appropriate statistical models to be used in our pipeline, we compared GSVA scores derived from gene expression data to predicted scores from random forest and XGBoost models with and without gene filtering with LASSO. This comparison was made for all Hallmark pathways across one-fold of the pan-cancer dataset. Both random forest approaches slightly outperformed XGBoost, and while the random forest model preformed marginally better without the LASSO feature selection (0.5413 vs 0.5362), this difference was not significant and therefore we chose to include LASSO for its ability to aid in identifying important genes for pathway score predictions and to prevent overfitting to the training data **(Suppl. Fig. 1)**. LASSO regression models were trained to predict GSVA scores for each pathway using the R package *glmnet*. The input data, ***X***, was a concatenated dataset of the mutation and copy number data **(Fig. 1b)**.

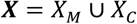

Where:

*X_M_* = {*discretized gene* − *level* somatic variants 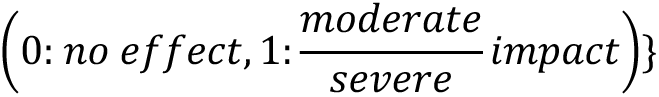

*X_C_* = {*discretized gene* − *level* copy number variations

(−2: *large deletion*, −1: *moderate deletion*, 0: *diploid*, 1: *moderate amplification*, 2: *large amplification*)}

The optimal lambda value was determined based on 5-fold cross validation and used to train a final LASSO model. The list of genes with coefficients greater than zero was collected for each pathway model. Separate models were additionally trained based only on the subset of genes included in the 410 MSK-IMPACT targeted sequencing panel.

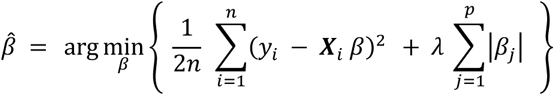

Where:

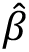 = *Estimated coefficients*

*y_i_* = *Gene set enric*ℎ*ment scores* (*from Eq. 1) for sample i*

***X****_i_* = *Integrated set of mutations and CNVs for sample i*

*λ* = *Regularization parameter*

|*β*_j_| = *Absolute value of t*ℎ*e coefficients* (*L*1 − *penalty*) *for gene j*

For each pathway, the concatenated mutation and copy number dataset was filtered to only include the subset of genes identified from the corresponding LASSO model. This filtered dataset, *X*^’^, consisting of a subset of mutations and CNAs with non-zero coefficients from the LASSO model wasthen used to train a random forest model to predict gene set enrichment score of signatures *ŷ* across all samples **(Fig. 1b)**.

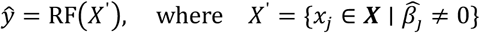

Random forest regression models were trained using the R package *randomForest*^66^. Each model used 100 trees and an mtry value equal to the square root of the number of input variables. Pathway models were trained and evaluated based on 5-fold cross validation. Correlation of real vs predicted pathway scores from out-of-fold test data was determined based on Pearson Correlation. UMAP visualization of individual cancer types was based on genomic-based pathway scores from out-of-fold test and validation datasets using the R package *umap*. Principal component analysis was used to compare disease subtypes in breast, lung, and brain cancers **(Fig. 1c)**. Biplots and component loadings were visualized using the R package *PCAtools*.

### Pathway phenotype grouping and classification

MSigDB Hallmark, KEGG (Legacy), Reactome, and ImmuneSigDB pathway collections were grouped into 11 pathway phenotypes modeled after known cancer hallmarks (DNA repair, evasion of growth suppressors, hormone signaling, avoiding immune destruction, tumor promoting inflammation, metabolism, activating invasion & metastasis, sustained proliferative signaling, unlocking phenotypic plasticity, resistance of cell death, and inducing angiogenesis pathways). Pathways related to each phenotype were extracted using keywords identified through literature search **(Suppl. Table 1)**. Immune pathways included the top 100 ImmuneSigDB pathway with highest correlation between RNA-based and genomic-based pathway scores in addition to alternate pathways from other pathway collections based on key words.

### PhenoSurv

Our model PhenoSurv takes input genomic-based pathway scores and generates phenotype-specific latent dimensions which are prospectively used to predict disease-free or progression-free survival **(Fig. 1d)**. The model architecture and training were done using *PyTorch*^83^ (version 2.4.1). Single sample pathway scores are grouped into phenotype classifications as specified above.

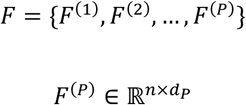

Where,

*F*^(*P*)^ = *pat*ℎ*way score matrix for p*ℎ*enotypic group P*

*n* = *number of samples*

*d_P_* = *number of feautures* (*pat*ℎ*way scores*) *in p*ℎ*enotypic group P*

The encoder uses linear layers with leaky ReLU activation functions to reduce the dimensionality of each phenotype group down to a single dimension.

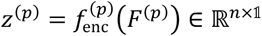

Where,

*f*^(*p*)^ = *encoderfunction*(*linearlayerswit*ℎ*leakyReLU*)

*z*^(*p*)^ = *latent representation of p*ℎ*enotype p*

The single latent dimensions from all phenotypes are then concatenated into *Z*.

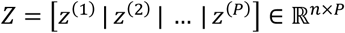

This latent space representation of the phenotypes is modeled using a Cox survival model, implemented in the package *TorchSurv*^84^.

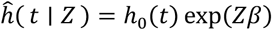

Where,

*ĥ*(*t* ∣ *Z*) = ℎ*azard function from Cox model*

ℎ_0_(*t*) = *baseline* ℎ*azard*

*β* = *coefficients of t*ℎ*e Cox model*

Concurrently, the decoder reconstructs each phenotype group from the latent dimensions.

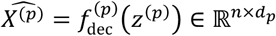

Where,

*f*^(*p*)^ = *decoder function for p*ℎ*enotype p*

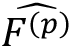 = *reconstructed score for p*ℎ*enotype p*

During model training, the total loss is represented as a sum of the negative partial log likelihood from the Cox survival model, the Kullback-Leibler divergence, and the binary cross entropy based on each phenotype reconstruction. The binary cross entropy for each phenotype is weighted based on the number of datapoints reconstructed. The negative partial log-likelihood from the cox model is weighted based on the number of events. The loss function for the PhenoSurv is shown below:

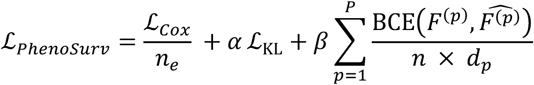

Where,

ℒ*_Cox_* = *Negative partial log* − *likeli*ℎ*ood from t*ℎ*e Cox model*

ℒ_KL_ = *Kullback* − *Leibler divergence*

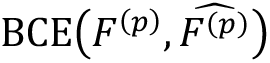 = *Binary cross*

− *entropy loss between orignial input and reconstructed p*ℎ*enotype scores*

*α, β* = ℎ*yperparameters controling contributions of KL divergence and reconstruction loss*

*n* = *number of samples*

*n_e_* = *number of events*

*d_P_* = *number of features* (*pat*ℎ*way scores*) *in p*ℎ*enotypic group P*

Models were trained using Adam optimization to determine optimal hyperparameter settings for each cohort. Concordance index from final model predictions was measured based on 5-fold cross validation of each dataset. Impact on model performance was measured based on weight decay [0, 0.0001, 0.001], batch size [8, 16, 32], number of epochs [50, 100, 1000, 2000], learning rate [0.01, 0.001, 0.0001], and dropout [0, 0.2].

To interpret the relevance of individual pathways within the phenotype groups of our models, Shapley values were calculated using DeepSHAP^85^. Trained PhenoSurv models for each dataset were used to calculate Shapley values of each pathway using the DeepExplainer function from the Python package *shap* **(Fig. 1e)**.

### Comparison to related survival models

To assess the performance of PhenoSurv, we tested two highly utilized survival models: random survival forest^86^ and DeepSurv^87^. These comparisons were performed using either the original genomic data as input or using the genomic-based pathway scores imputed using PhenoMap. Using the R package *randomForestSRC*, the performance of random survival forest model was evaluated on the TCGA and MSK datasets using either disease-free or progression-free survival and 5-fold cross validation. For each fold, optimal mtry and node size hyperparameters were determined using the tune function. DeepSurv models were trained on the same 5 cross-validation folds for each dataset using the R package *mlr3*^88^. Hyperparameters for the DeepSurv models were tuned using Adam optimization based on dropout [0 – 1], weight decay [0 – 0.5], learning rate [0 – 1], and number of nodes [0 – 16]. Concordance index of the test datasets based on the final optimal model was measured using the R package *survcomp*^89^. To identify significant differences between the three model performances, ANOVA analysis was performed to identify differences across models, followed by post-hoc Tukey HSD for pairwise comparison. To assess the reproducibility of the final PhenoSurv and random forest models, feature importance was first calculated based on either mean SHAP values (PhenoSurv) or Gini index (random survival forest). Across the 5 trained models for each training fold, the Spearman correlation was calculated. Mean Spearman correlations were used to compare model reproducibility.

### Computational analysis & data visualization

All dataset curation, genomic-based pathway scoring, and data visualization were performed in R version 4.4.277. Deep learning survival analysis was performed in Python version 3.11. R packages *ggplot2, ggpubr, survival* and *survminer,* were used for data visualization. For pathway biomarker discovery, pathway scores were split based on maximally-ranked statistics from the TCGA datasets. Validation in the MSK datasets was done using the same cutpoint identified from TCGA to split high-and low-signaling groups.

## Data and code availability

All data used for this study came from publicly available sources, including TCGA pan-cancer data from NCI Genomic Data Commons (https://gdc.cancer.gov/), and CPTAC and MSK datasets from cBioPortal (https://www.cbioportal.org/datasets). Pretrained PhenoMap models for over 1,500 pathways with robust performance in validation data across breast, lung, and brain cancers are available on The Open Science Framework (https://osf.io/94e32/). Code for PhenoMap and PhenoSurv models, downstream data analyses, and figure generation is available on Github (https://github.com/nathlab-coh/PhenoMap-PhenoSurv).

