## Supplementary Information for "Phenotypic inference from sparse tumor genomes informs an explainable deep-learning model for cancer prognosis"

**Supplementary Information: Transforming Genomic Profiles to Phenotypic Insights to Advance Explainable Deep Learning for Cancer Prognosis**

**SUPPLEMENTARY FIGURES**

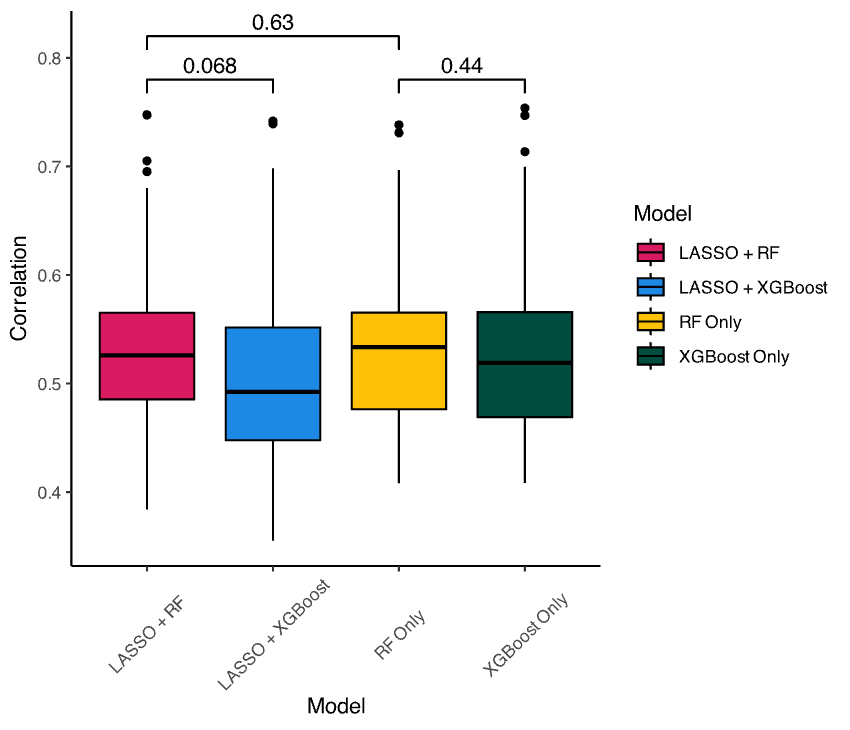

**Supplementary Figure 1.** GSVA vs genomics-based model calculations. Boxplots comparing correlations between gene expression-based pathway enrichment scores vs. predicted pathway scores from genomics-based models. Different combinations of modeling approaches included prior feature selection with LASSO in combination with random forest (RF) or XGBoost models, or using RF/XGBoost models with all available features.

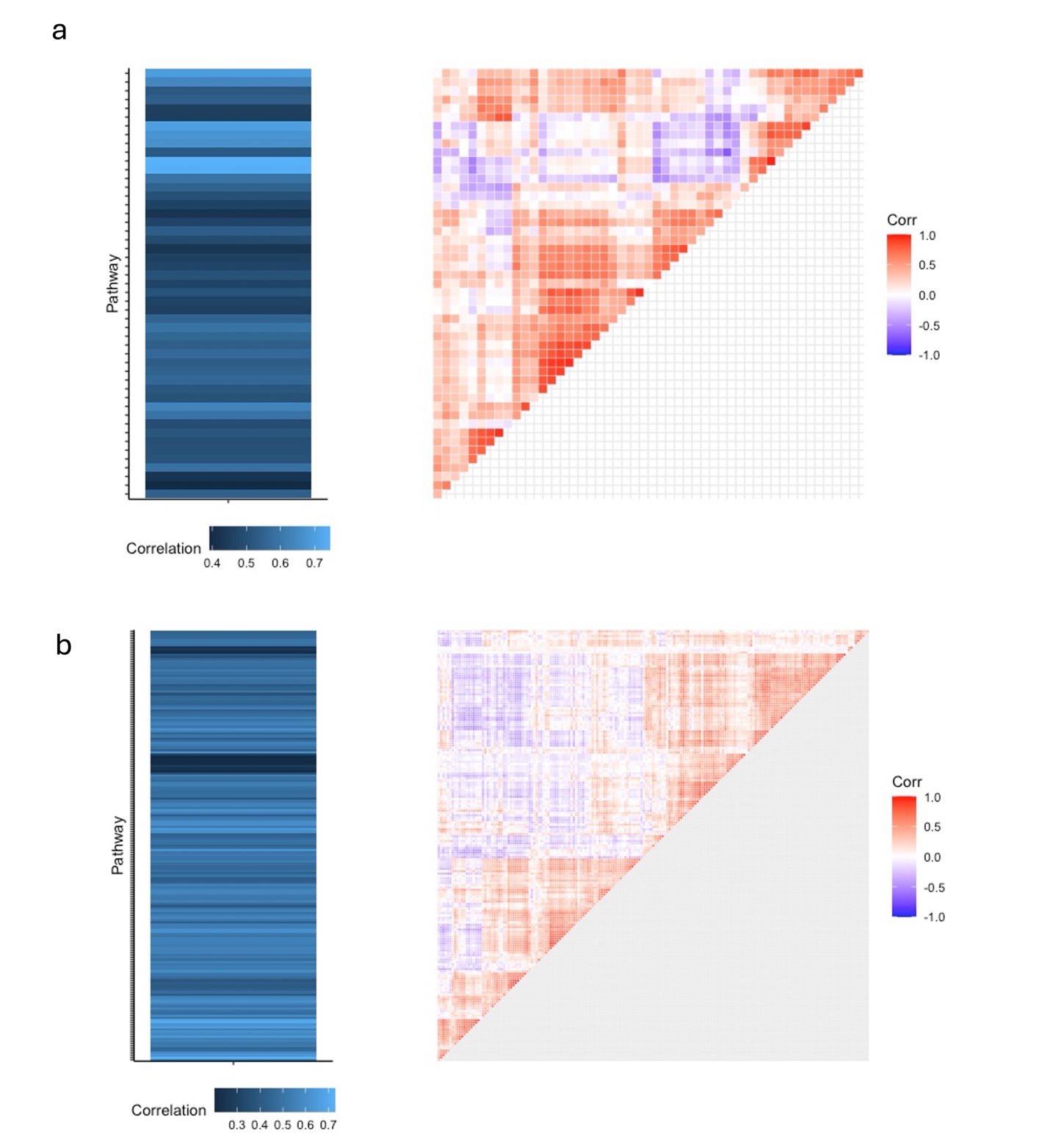

**Supplementary Figure 2**. Correlation matrix and hierarchical clustering of real pathways scores (right) and correlation of real vs predicted scores of each pathway (left) for a) hallmark and b) KEGG pathways

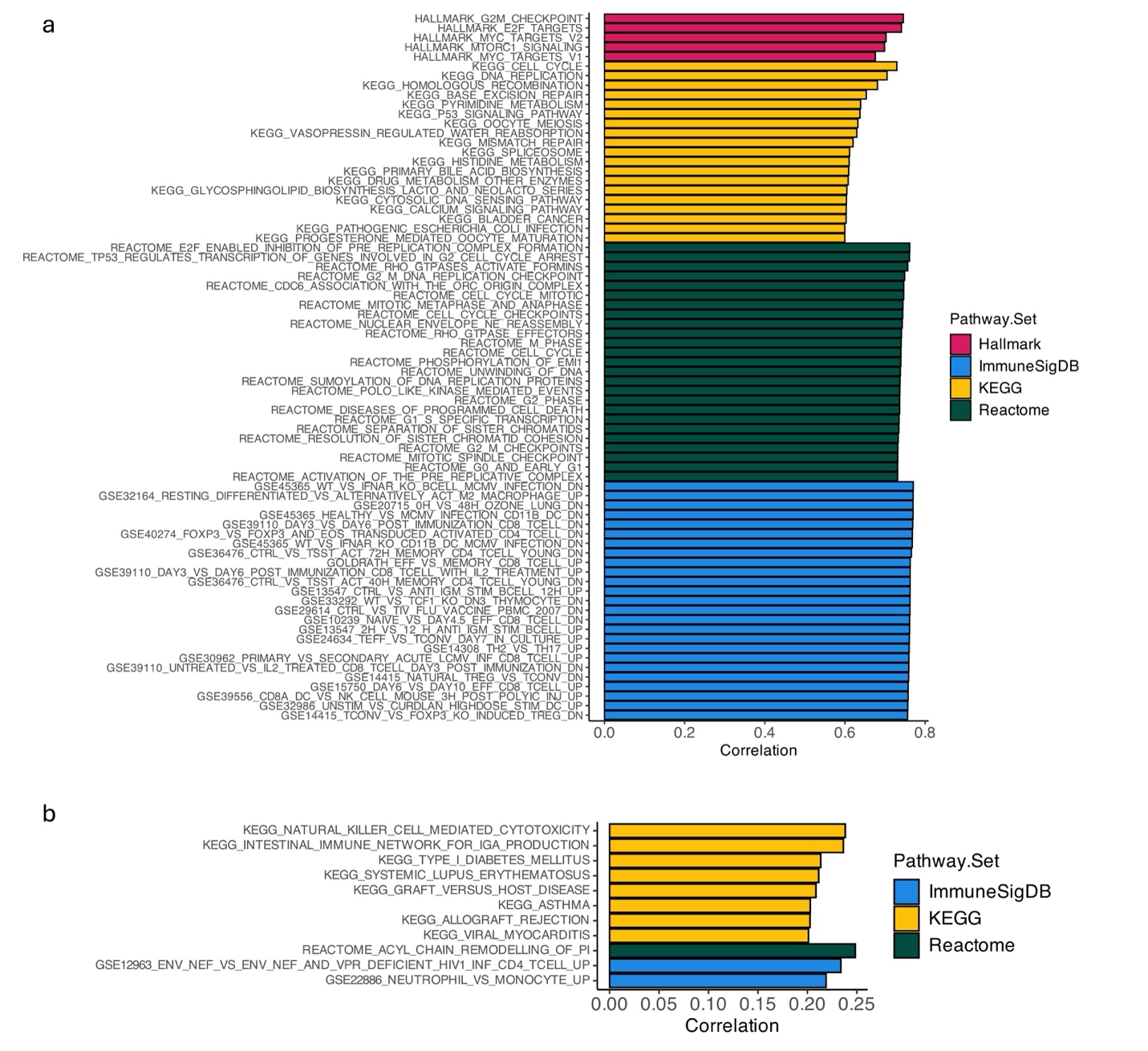

**Supplementary Figure 3.** Correlation between PhenoMap pathway scores vs GSVA pathway scores in TCGA pan-cancer out-of-fold test data. a) Pathways in top 10% of pathway set (Hallmark, KEGG), or top 25 pathways (Reactome, ImmuneSigDB). b) Pathways with correlation below 0.25.

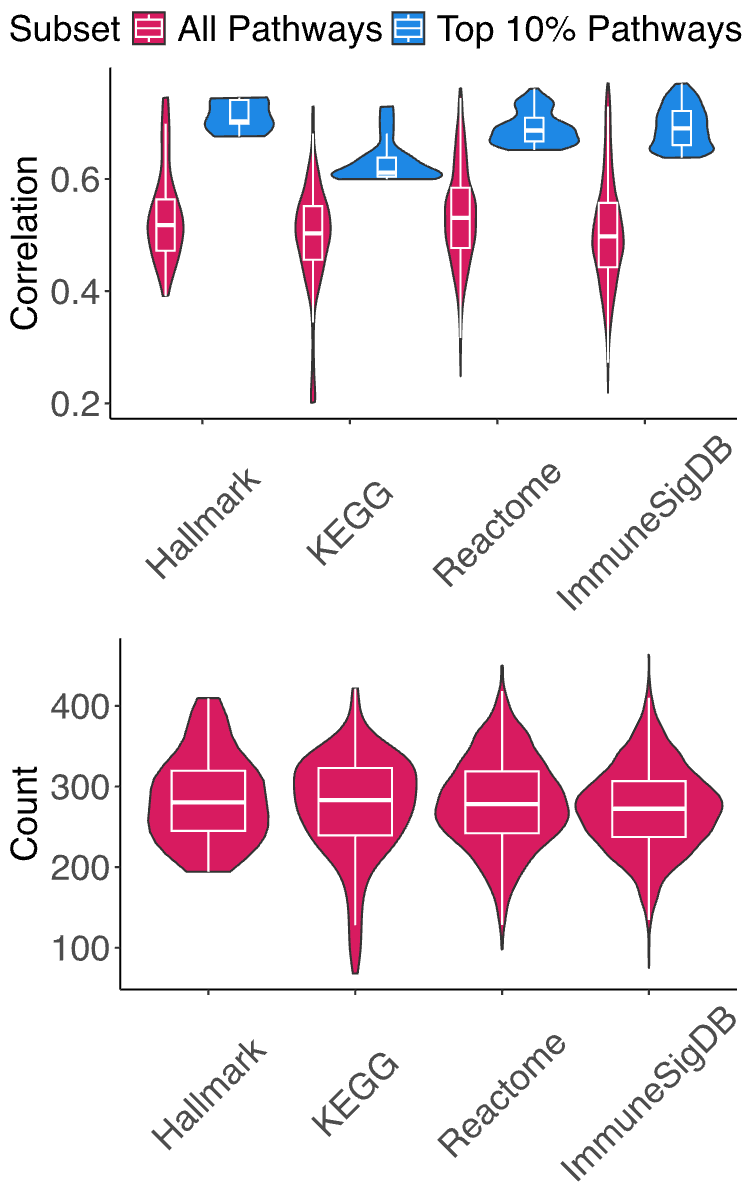

**Supplementary Figure 4.** Violin plot displaying the number of gene features selected by LASSO in different classes of pathway prediction models.

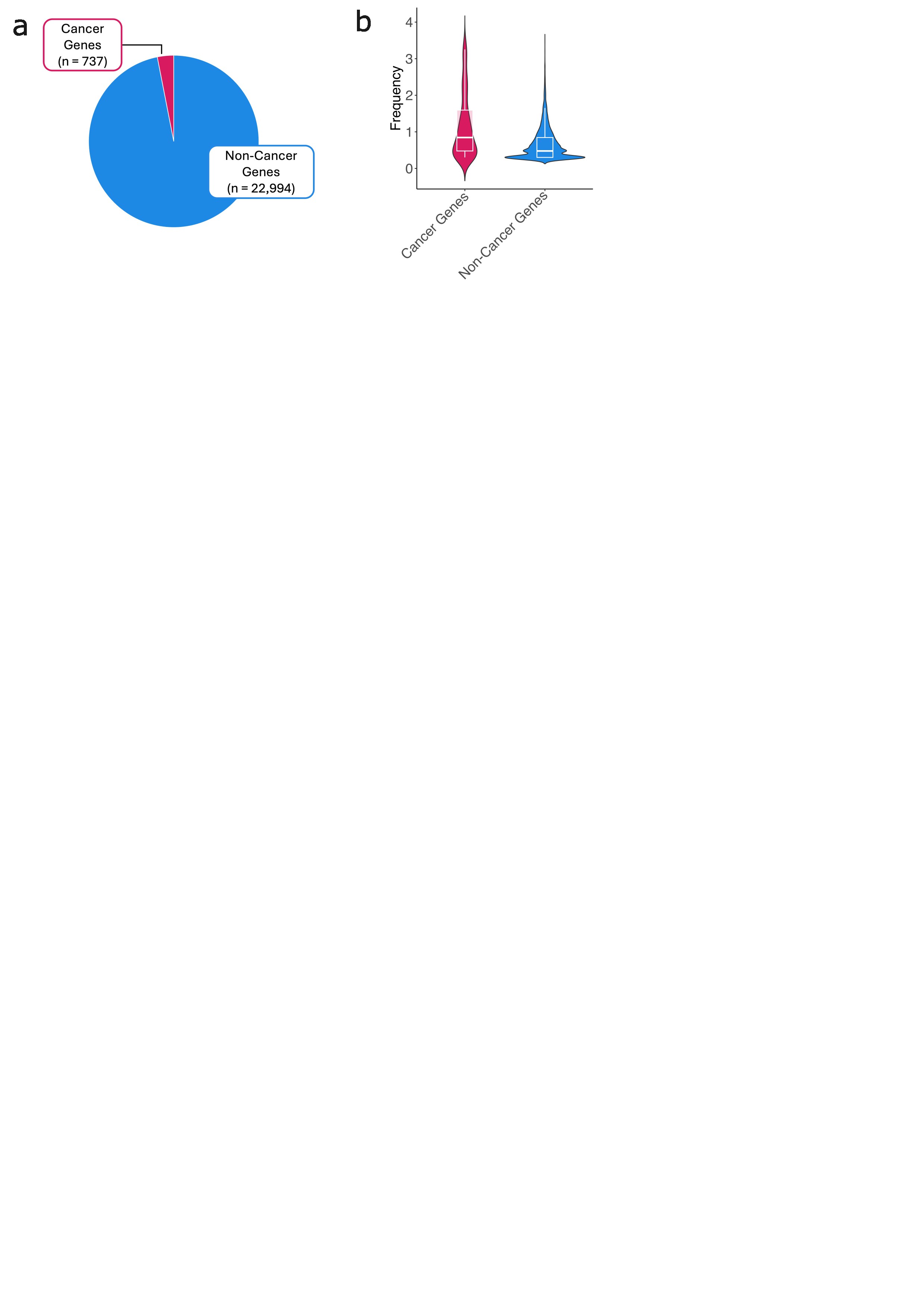

**Supplementary Figure 5.** a) Pie chart showing proportion of cancer genes selected by the LASSO models represented in COSMIC Cancer Gene Census vs. non-cancer genes. b) Violin plot showing frequency each COSMIC cancer gene or non-cancer genes was selected by LASSO models.

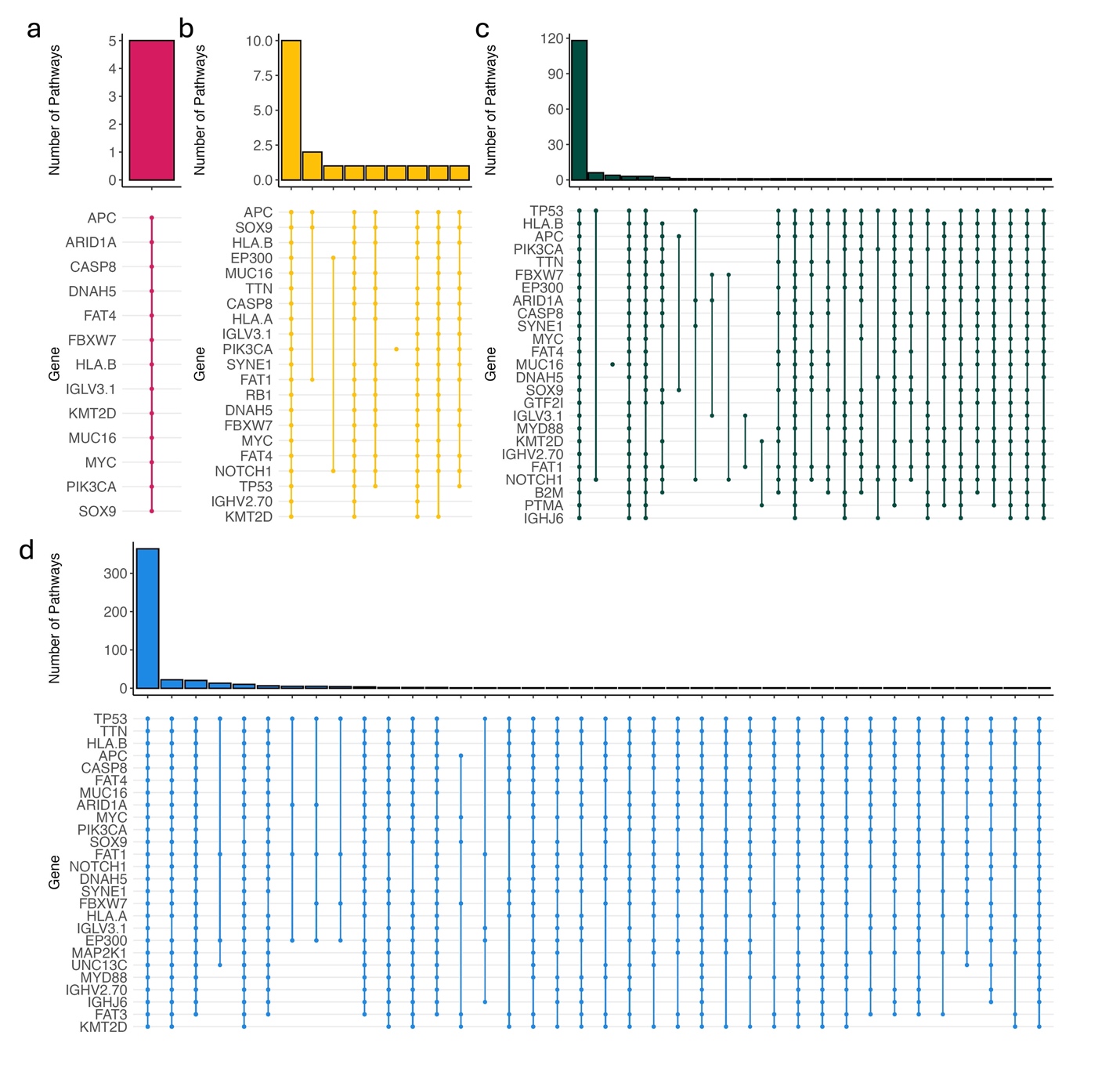

**Supplementary Figure 6.** Combinations of top 1% most frequent genes chosen by LASSO model for prediction of a) Hallmark, b) KEGG, c) Reactome, and d) ImmuneSigDB top 10% best predicted pathways.

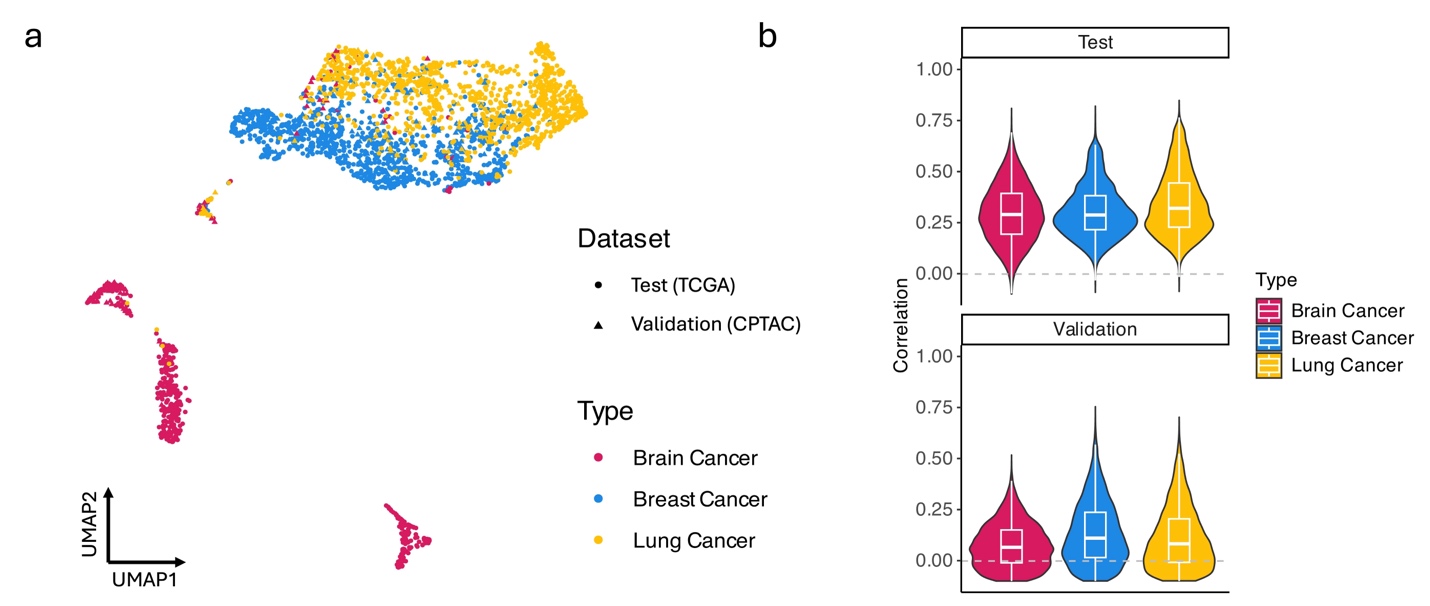

**Supplementary Figure 7.** Validation of genomic-based pathway scores. a) UMAP projections of TCGA test samples (breast n = 981, lung n = 967, and brain 505) and CPTAC validation samples (breast n = 121, lung n = 109, and brain n = 95 cancers) based on genomic-based pathways grouped by cancer type. b) Violin plots comparing correlation of genomic-based and RNA-based pathway scores by cancer type in TCGA test and CPTAC validation cohorts.

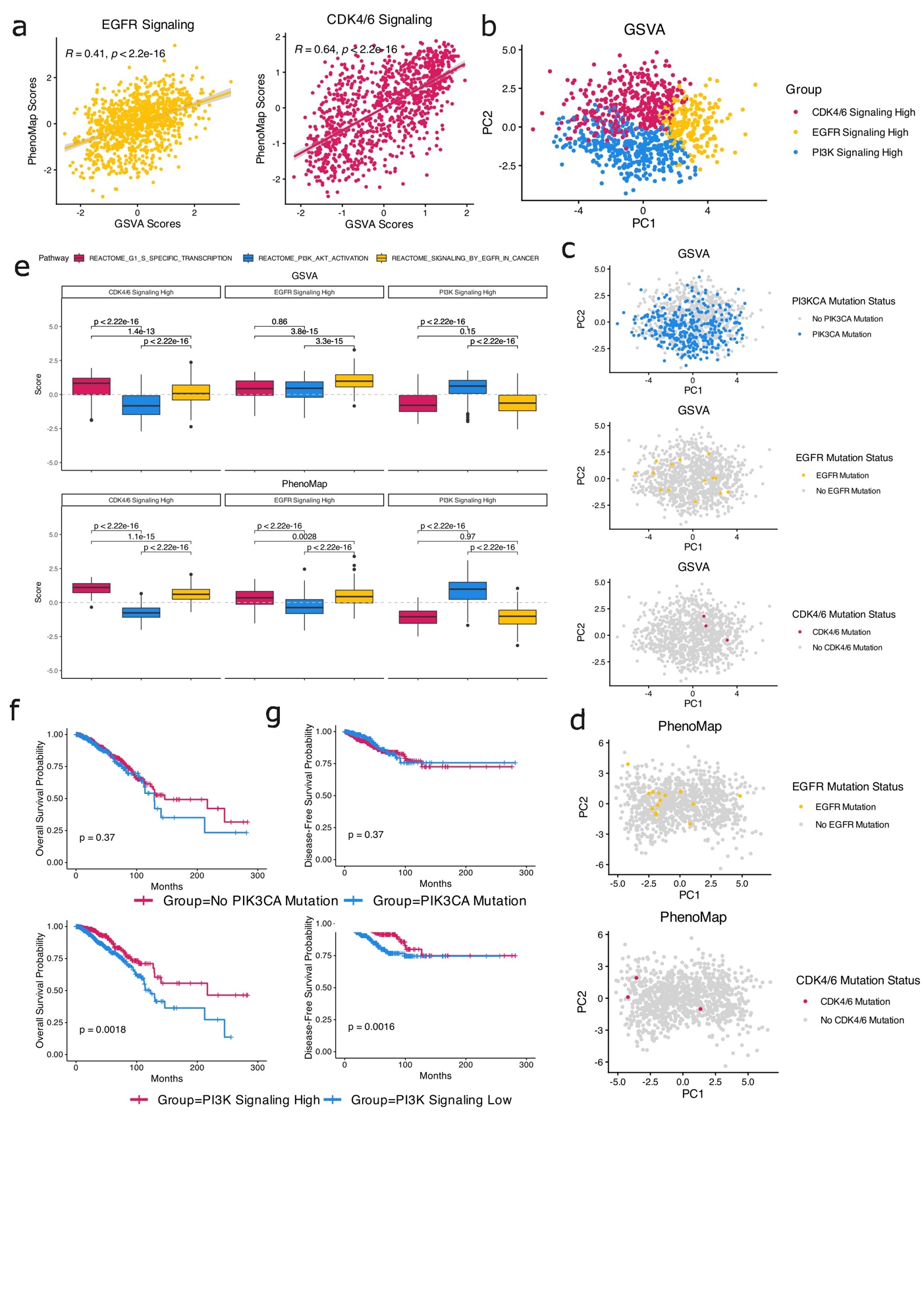

**Supplementary Figure 8.** a) Dot plot and spearman correlation of GSVA and PhenoMap pathway scores in breast cancer. b) PCA projections of breast cancer clusters based on most activated signaling pathway. Clusters from GSVA scores were EGFR signaling high (n = 466), CDK4/6 signaling high (n = 206), and PI3K signaling high (n = 419). Clusters from PhenoMap scores were EGFR signaling high (n = 206), CDK4/6 signaling high (n = 197), and PI3K signaling high (n = 331). c) PCA projections based on GSVA scores relative to gene mutation status. d) PCA projections based on GSVA scores relative to gene mutation status. e) Boxplots comparing GSVA and PhenoMap scores of each cluster based on representative signaling pathways in breast cancer. f) Overall survival in breast cancer cases with or without PIK3CA mutation (top) or in breast cancer cases with PhenoMap scores above or below mean value in the Reactome PI3K AKT activation pathway (bottom). g) Disease-free survival in breast cancer cases with or without PIK3CA mutation (top) or in breast cancer cases with PhenoMap scores above or below mean value in the Reactome PI3K AKT activation pathway (bottom).

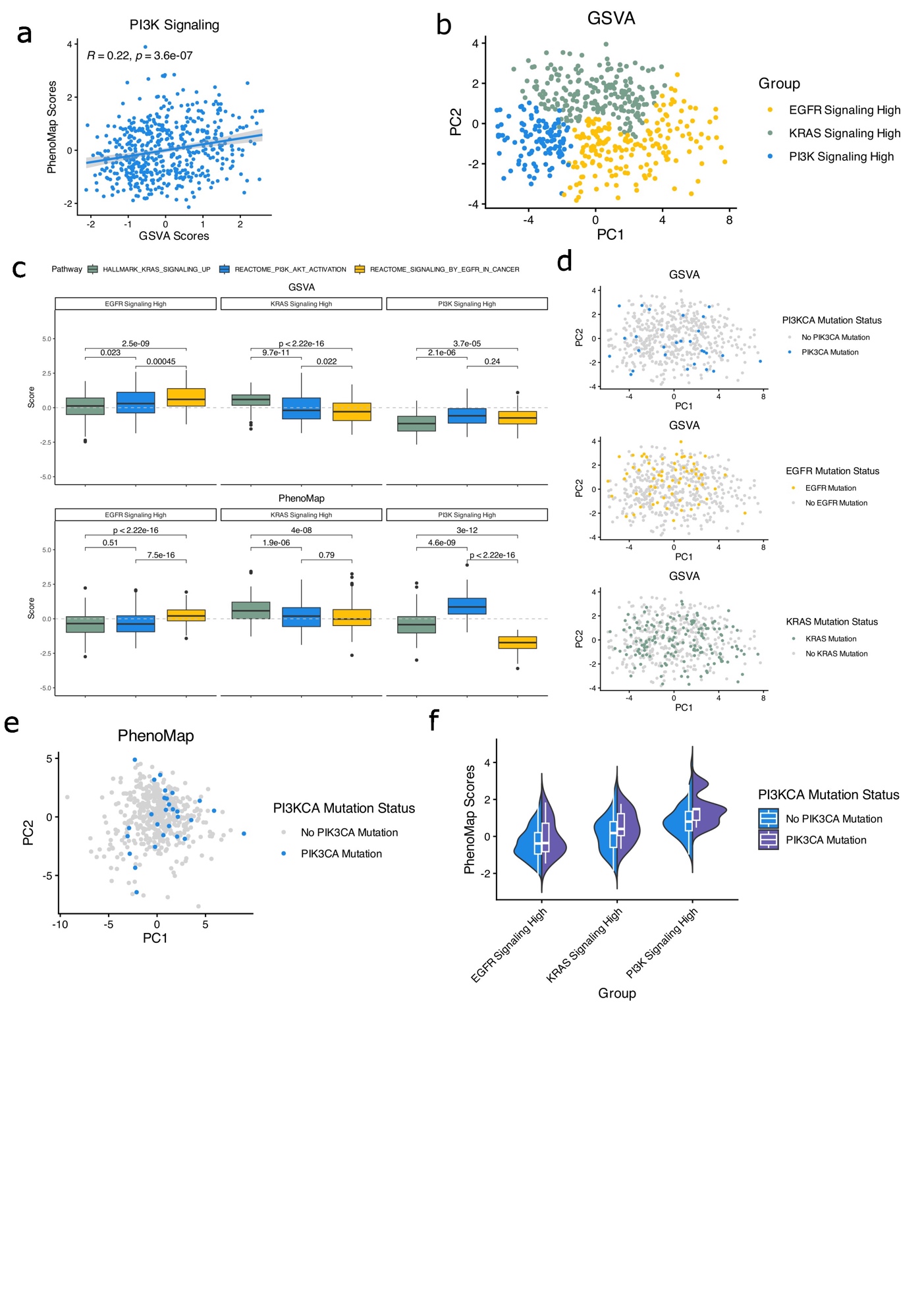

**Supplementary Figure 9.** a) Dot plot and spearman correlation of GSVA and PhenoMap pathway scores in lung cancer. b) PCA projections of lung cancer clusters based on most activated signaling pathway. Clusters from GSVA scores were EGFR signaling high (n = 185), KRAS signaling high (n = 207), and PI3K signaling high (n = 111). Clusters from PhenoMap scores were EGFR signaling high (n = 255), KRAS signaling high (n = 196), and PI3K signaling high (n = 52). c) Boxplots comparing GSVA and PhenoMap scores of each cluster based on representative signaling pathways in lung cancer. d) PCA projections based on GSVA scores relative to gene mutation status. e) PCA projections based on GSVA scores relative to gene mutation status. f) Split violin plot of PhenoMap PI3K Signaling scores by cluster in patients with or without PI3KCA mutation.

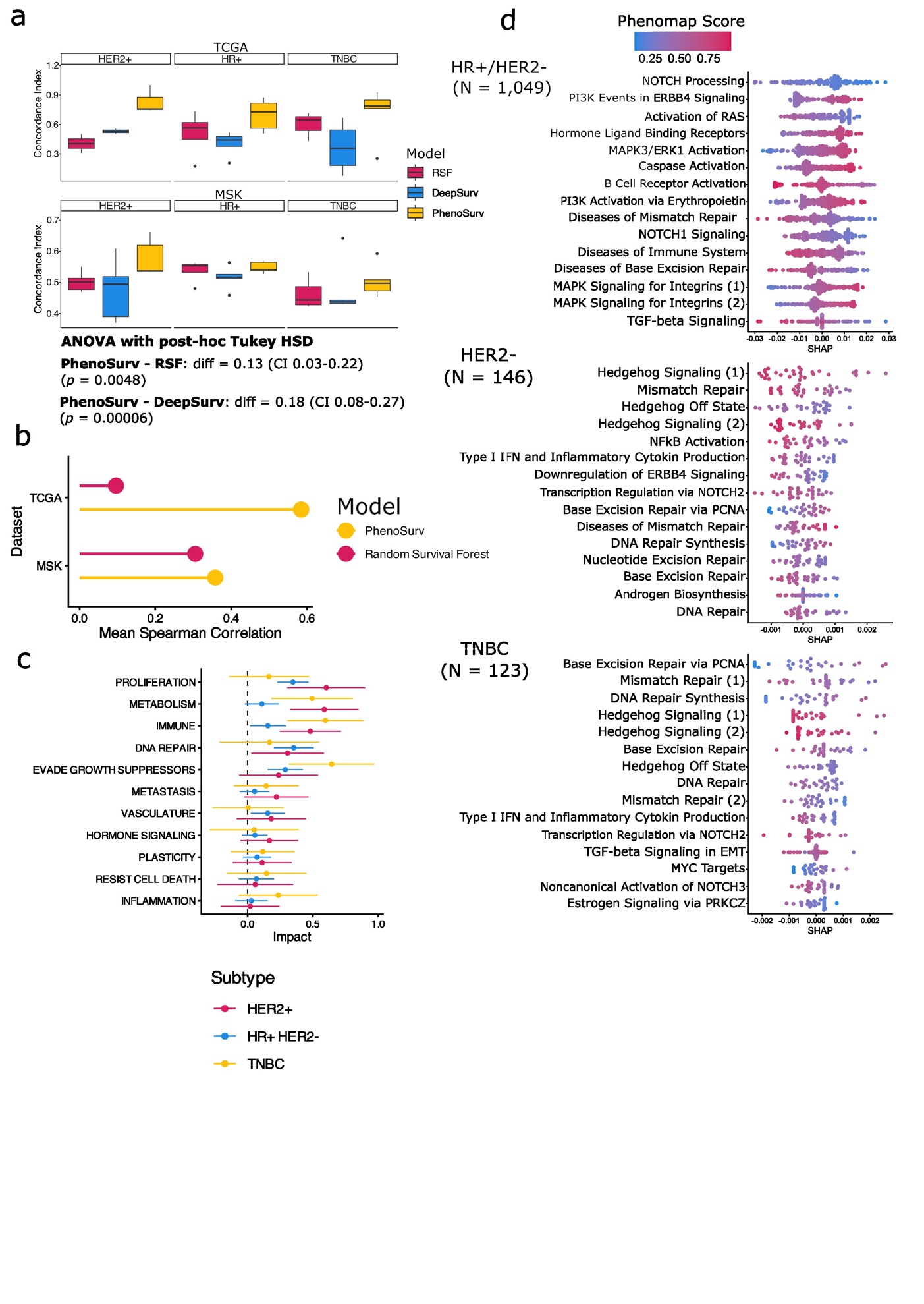

**Supplementary Figure 10.** a) Boxplot comparing concordance indices of PhenoSurv and previously published models including Random Survival Forests and DeepSurv using expression-based pathway scores and genomic data as input by subtype and dataset in breast cancer. b) Mean spearman correlation of feature importance between five dataset splits based on either mean SHAP values (PhenoSurv) or Gini index (RSF). c) Comparison of hazard ratios of phenotype-based latent dimensions from trained PhenoSurv models in MSK breast cancer dataset. d) Beeswarm plots of top 15 pathways with highest SHAP values from PhenoSurv models in MSK breast cancer dataset.

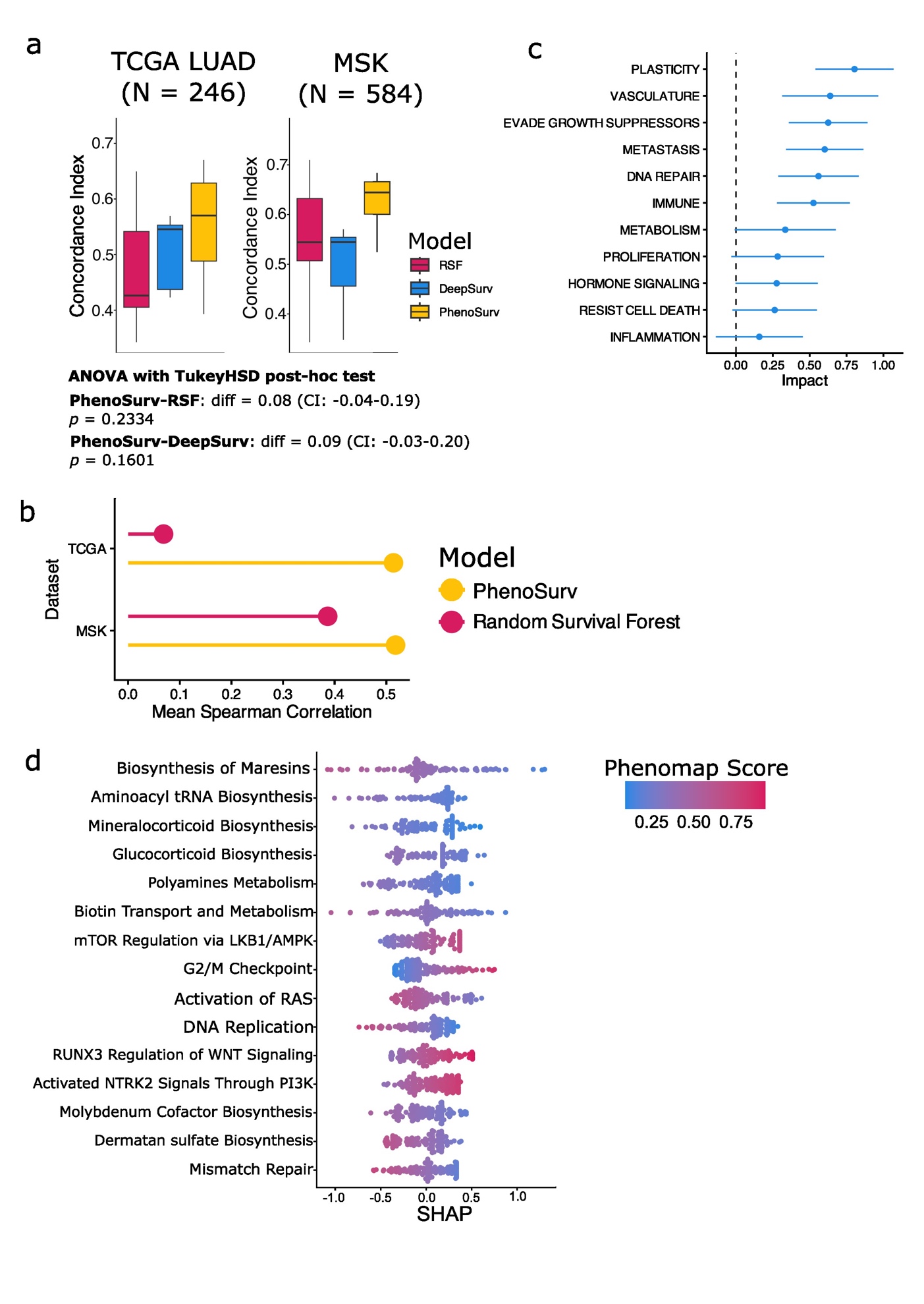

**Supplementary Figure 11.** a) Boxplot comparing concordance indices of PhenoSurv and previously published models including Random Survival Forests and DeepSurv using expression-based pathway scores and genomic data in MSK lung cancer. b) Mean spearman correlation of feature importance between five dataset splits based on either mean SHAP values (PhenoSurv) or Gini index (RSF). c) Comparison of hazard ratios of phenotype-based latent dimensions from trained PhenoSurv models in MSK lung cancer dataset. d) Beeswarm plots of top 15 pathways with highest SHAP values from PhenoSurv models in MSK lung cancer dataset.

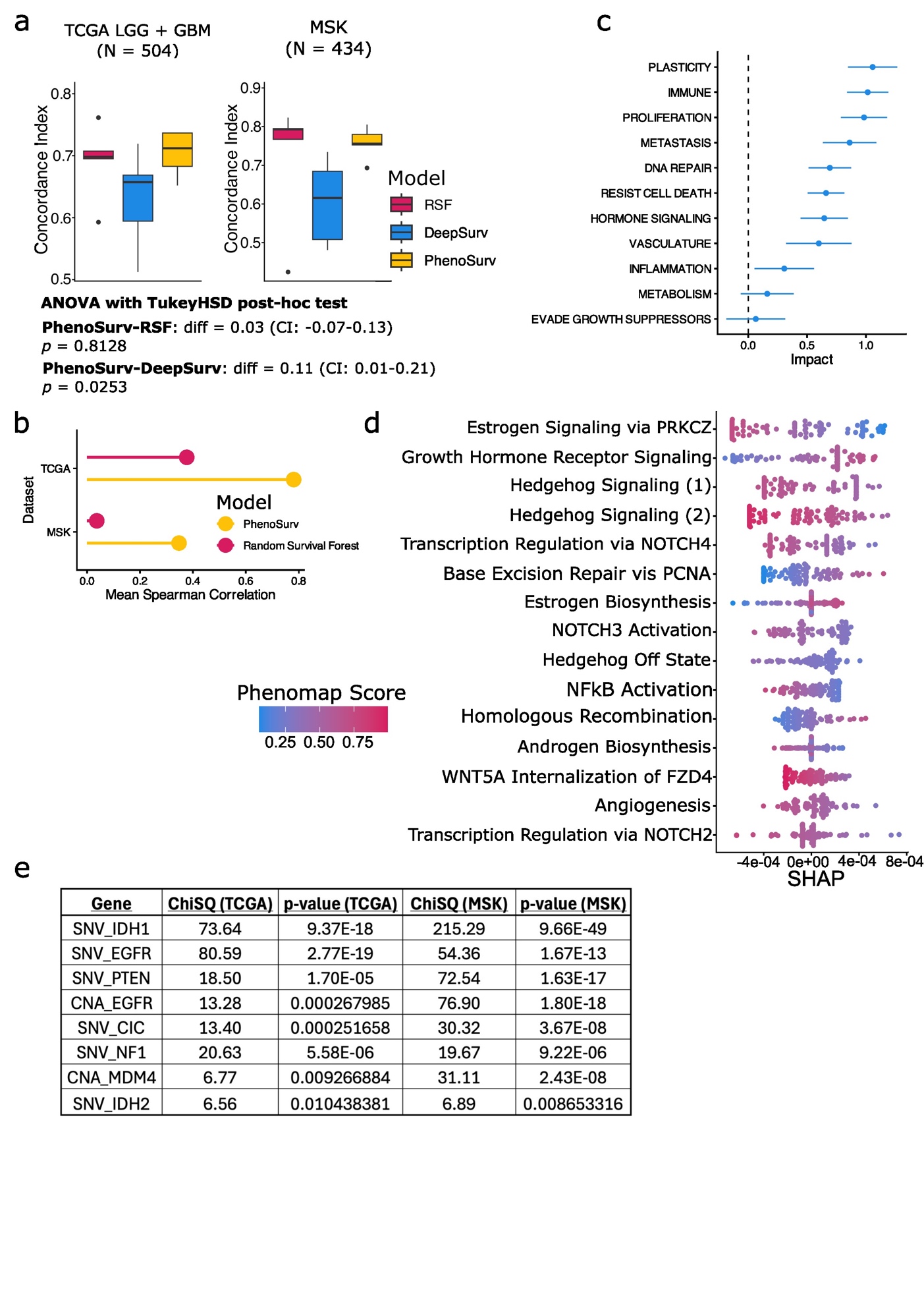

**Supplementary Figure 12.** a) Boxplot comparing concordance indices of PhenoSurv and previously published models including Random Survival Forests and DeepSurv using expression-based pathway scores and genomic data in MSK brain cancer dataset. b) Mean spearman correlation of feature importance between five dataset splits based on either mean SHAP values (PhenoSurv) or Gini index (RSF). c) Comparison of hazard ratios of phenotype-based latent dimensions from trained PhenoSurv models in MSK brain cancer dataset. d) Beeswarm plots of top 15 pathways with highest SHAP values from PhenoSurv models in MSK breast cancer dataset. e) List of significant gene biomarkers identified by PhenoSurv based on progression-free survival in brain cancer datasets.

**SUPPLEMENTARY TABLES**

**Supplementary Table 2.** Key words used to extract list of pathways for model phenotypes

| **Phenotype** | **Alias** | **Key words** |
| --- | --- | --- |
| Genome instability and mutation | DNA REPAIR | “DNA_REPAIR”, “BASE_EXCISION_REPAIR”, “HOMOLOGOUS_RECOMBINATION”, “MISMATCH_REPAIR”, “NUCLEOTIDE_EXCISION_REPAIR” |
| Evasion of growth suppressors | EVADE GROWTH SUPPRESSORS | “G2M”, “G2”, “G1”, “_S_”, “P53”, “CELL_CYCLE”, “AUTOPHAGY”, “CHECKPOINT” |
| Hormone signaling | HORMONE SIGNALING | “ANDROGEN”, “ESTROGEN”, “ESR”, “HORMONE”, “PROGESTERONE” |
| Avoiding immune destruction | IMMUNE | “IMMUNE”, “B_CELL”, “T_CELL”, “NK_CELL” |
| Tumor promoting inflammation | INFLAMMATION | “IL2”, “IL6”, “INFLAMMATORY”, “INTERFERON”, “TNFA”, “JAK”, “STAT”, “TOLL_LIKE”, “TLR”, “NOD_LIKE”, “NLR”, “CHEMOKINE”, “CYTOKINE” |
| Cancer metabolism | METABOLISM | “METABOLISM”, “BIOSYNTHESIS”, “OXIDATIVE_PHOSPHORYLATION”, “PENTOSE_PHOSPHATE_PATHWAY”, “CITRATE_CYCLE”, “REACTIVE_OXYGEN_SPECIES”, “GLYCOLYSIS” |
| Activating invasion and metastasis | METASTASIS | “TGF_BETA”, “HEDGEHOG”, “METASTASIS” |
| Sustained proliferative signaling | PROLIFERATION | “KRAS”, “MTOR”, “TGF”, “PI3K”, “AKT”, “MYC”, “E2F”, “MEK”, “ERK”, “ERBB”, “MAPK”, “PROLIFERATION” |
| Unlocking phenotypic plasticity | PLASTICITY | “WNT”, “EMT”, “EPITHELIAL_MESENCYMAL_TRANSITION”, “PLASTICITY” |
| Resisting cell death | RESIST CELL DEATH | “APOPTOSIS”, “BCL2”, “CASPASE”, and “BAX” |
| Inducing angiogenesis | VASCULATURE | “ANGIOGENSIS”, “NOTCH”, “VEGF”, “VASCULATURE” |

**Supplementary Table 1.** Correlation of pathway scores from GSVA vs scores predicted by PhenoMap in TCGA pan-cancer dataset including 6,722 tested pathways from Hallmark, KEGG, Reactome, and ImmuneSigDB pathway sets.

**Supplementary Table 3.** Concordance indices of all cancer types for PhenoSurv, Random Survival Forest, and DeepSurv.

**Supplementary Table 4.** List of significant pathway biomarkers identified from brain cancer PhenoSurv model.
